# Frequent Non-random Shifts in the Temporal Sequence of Developmental Landmark Events during Teleost Evolutionary Diversification

**DOI:** 10.1101/239954

**Authors:** Fumihiro Ito, Tomotaka Matsumoto, Tatsumi Hirata

## Abstract

Morphological transformations can be generated by evolutionary changes in the sequence of developmental events. In this study, we examined the evolutionary dynamics of the developmental sequence on a macroevolutionary scale using the teleost. Using the information from previous reports describing the development of 31 species, we extracted the developmental sequences of 19 landmark events involving the formation of phylogenetically conserved body parts; we then inferred ancestral developmental sequences by two different parsimony-based methods—event-pairing (Jeffery, Bininda-Emonds, Coates & Richardson, 2002a) and continuous analysis (Germain & Laurin, 2009). The phylogenetic comparisons of these sequences revealed event-dependent heterogeneity in the frequency of sequence changes. Most of the sequence changes occurred as exchanges of temporally neighboring events. The phylogenetic analyses suggested that the ancestral species had experienced frequent changes in developmental sequences. Although the analyses showed that these heterochronic changes accumulated along phylogenic time, the precise distribution of the changes over the teleost phylogeny remains unclear due to technical limitations. Finally, this first comprehensive analysis of teleost developmental sequences will provide solid ground on which to elucidate the significance of developmental timing in animal morphological diversification.

## 1. Introduction

Development of multicellular organisms is characterized as a series of morphological events (Smith, 2001). Because individual events progress step by step in a hierarchical manner, one might assume that the temporal sequence of developmental events, i.e., the developmental sequence, does not readily change and is phylogenetically conserved among closely related species that share morphological characteristics. Consequently, if an evolutionary change occurs in the developmental sequence, it could have a significant impact on body patterning and lead to morphological diversity. Indeed, previous comparisons of developmental sequences have revealed rare epoch-making changes that can provide morphological uniqueness to one species (Strauss, 1990; Jeffery, Bininda-Emonds, Coates & Richardson, 2002a; Maxwell, Harrison & Lasson, 2010).

Another factor influencing the evolution of developmental sequences is the phylotypic period. This is the developmental period during which evolutionally distant animal species resemble each other. Historically, various models have proposed different time frames for the phylotypic period and discussed why evolutionary changes are difficult to occur in this period (Haeckel, 1874; Duboule, 1994; Raff, 1996; Richardson, 1999). Recently, quantitative approaches have begun to test these models. Transcriptome analyses showed that inter-species diversity is minimum during the middle phase of ontogenetic development (Domazet-Loso & Tautz, 2010; Kalinka et al., 2010; Irie & Kurtani, 2011), supporting the hourglass model that predicts that phenotypic diversity is large in early and late development, and restricted in between. In contrast, quantitative comparison of developmental sequences using 14 vertebrate and mammalian species indicated an opposite trend, namely that phenotypic variation between species is the highest in the middle phase of development (Bininda-Emond, Jeffery & Richardson, 2003). Together with some skepticism surrounding the actual existence of the conserved period (Poe & Wake, 2004; Richardson, 2012), the phylotypic period itself is still a controversial topic in evolutionary developmental biology and requires further rigorous testing with different datasets.

Despite the many stimulating theories and ongoing intensive discussion, there is little empirical evidence about evolutionary changes that may have occurred in the developmental sequences; few systematic comparisons have been made of the sequences of a wide range of developmental events covering the entire body in any class of animal. Therefore, there is little data available on how commonly or rarely the developmental sequences had actually changed during evolution. In recent decades, comparative methods to analyze developmental sequences have been developed by several groups (Nunn & Smith, 1998; Jeffery et al., 2002a; Jeffery, Richardson, Coates & Bininda-Emonds, 2002b; Jeffery, Bininda-Emonds, Coates & Richardson, 2005; Harrison & Larsson, 2008; Germain & Laurin, 2009). These methods compare the relative order of developmental events among different species and estimate potential evolutionary shifts of the events, i.e., “heterochronic shifts”, in a parsimonious manner (Schoch, 2006; Smirthwaitet, Rundle, Bininda-Emonds & Spicer, 2007; Sanchez-Villagra, Goswami, Weisbecker, Mock & Kuratani, 2008; Hautier et al., 2013; Laurin, 2014; Werneburg & Sanchez-Villagra 2015; Carril & Tambussi, 2017). Although most previous analyses focused on the developmental sequences for a specific organ or limited body part, we considered the methods themselves to be similarly applicable to a global analysis of developmental events comprising the entire body.

In this study, we conducted the first comprehensive survey of developmental sequences of teleosts. Teleosts comprise numerous species characterized by great morphological diversity (Nelson, Grande & Wilson, 2016) and shared conserved body structures including vertebrae, eyes, and paired fins (Romer & Parsons, 1986). Owing to the popularity of some teleosts as developmental research models, there are well-established staging tables for many teleost species covering common clear-cut developmental landmarks. Hence, the teleost can provide an ideal dataset for systematic analysis of developmental sequences. Among the widely used developmental landmarks, we chose 19 events that individually contribute to distinct body parts across the whole body. Using the dataset of 31 actinopterygian species, we compared the developmental sequences and reconstructed the ancestral sequences over the teleost phylogeny. Our analysis indicates that the developmental sequences have changed frequently during evolution. However, the changes were not random; the order of a few events seemed to have shifted more frequently than the others in the developmental sequence. These developmental sequence rearrangements may have contributed to shaping teleost morphological diversity.

## 2. Materials and Methods

### 2.1 Construction of the teleost phylogenetic tree

The overall topology of the phylogenetic tree followed the molecular phylogenetic relationship previously reported by Near et al. (2012, 2013). The minor branches missing in the tree were inserted based on the phylogenetic data obtained from Saitoh et al. (2011) and Yang et al. (2015) for Cypriniformes, Perez et al. (2007) and Friedman et al. (2013) for Cichliformes, and Pohl, Milvertz, Meyer & Vences (2015) for Cyprinodontiformes. The divergence times were determined using the public database TIMETREE, *The Timescale of Life* (Hedes & Kumar, 2009) (Supplemental file 1).

### 2.2 Data sampling

Information about developmental events was extracted from 31 published research articles that describe normal development of actinopterygian species (Figure 1). These developmental events were selected according to the following criteria: (1) clearly defined in most of the selected articles, (2) involved in a wide range of body parts and systems, and (3) occurring from early to late phases of embryogenesis. The 19 events examined in the study were the first instances of blood circulation (bc), caudal fin rays (cfr), eye pigmentation (ep), embryonic shield (es), the first somite (fs), hatching (h), heart beating/pulsing (hb), Kupffer’s vesicle (kv), lenses or lens placodes (le), a medial finfold (mff), a mouth opening (mo), olfactory vesicles/pits/placodes (olf), otoliths (oto), otic vesicles/placodes/primordia (ot), optic vesicles/placodes/primordia (op), pectoral fin buds (pfb), a tail bud (tb), three brain regionalization (tbr), and tail lift from the yolk (tl). The temporal order of the developmental events was ranked according to descriptions in the articles (Figure 1). When an article did not describe a developmental event, the event was treated as a missing datum.

**FIGURE 1.**
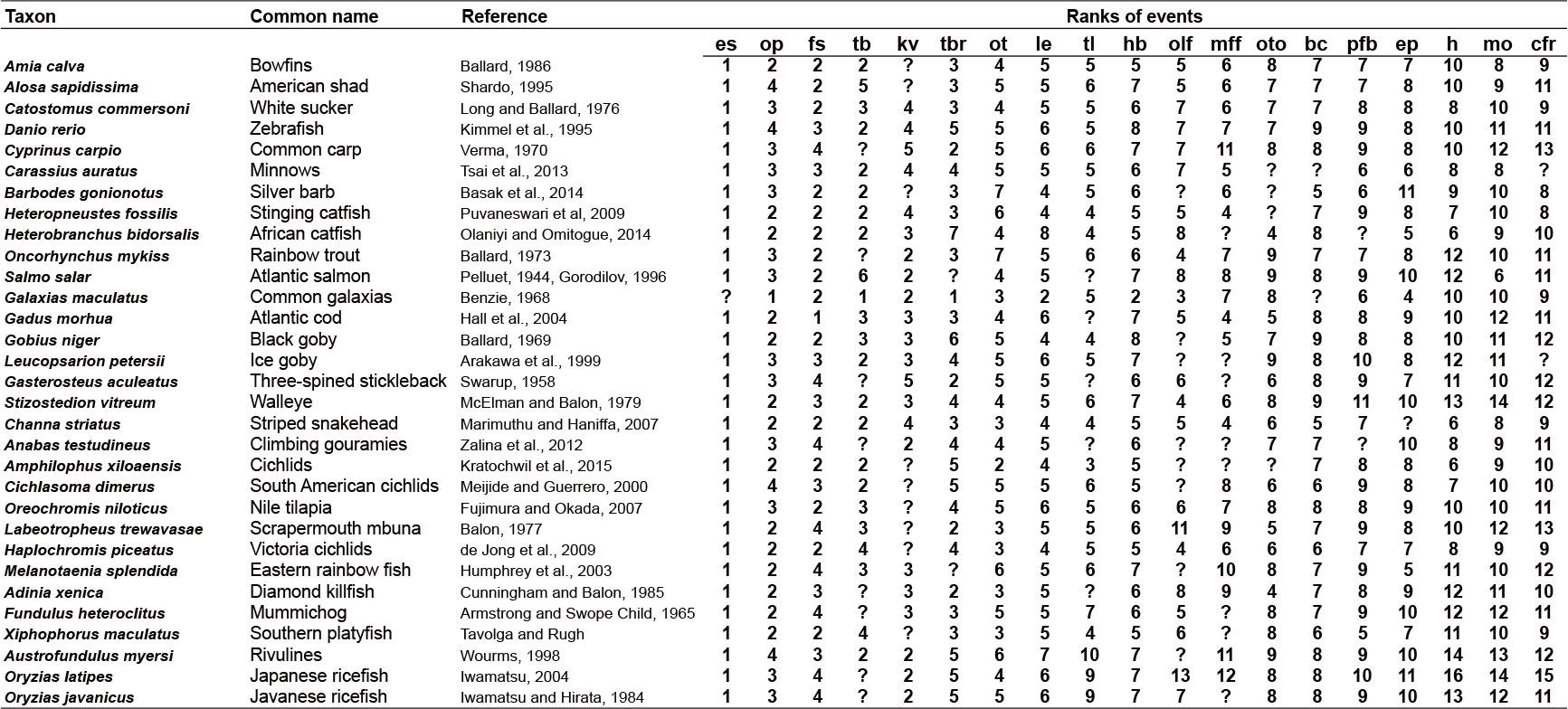
Temporal orders of developmental events in the 31 actinopterygian species. The temporal sequence of developmental events was extracted from the reference listed for each species. Abbreviations of developmental events are: bc, blood circulation; cfr, caudal fin rays; ep, eye pigmentation; es, embryonic shield; fs, first somite; h, hatching; hb, heart beating/pulsing; kv, Kupffer’s vesicle; le, lenses or lens placodes; mff, medial finfold; mo, mouth opening; olf, olfactory vesicles/pits/placodes; oto, otoliths; ot, otic vesicles / placodes/primordia; op, optic vesicles/placodes/primordia; pf, pectoral fin bud; tb, tail bud; tbr, three brain regionalization; and tl, tail lift from the yolk. The ranks of missing data are marked by ?.

### 2.3 Relative scaling of the ranks of developmental events

The raw ranks of individual developmental events were determined from the developmental sequences of extant species (Figure 1). When multiple events were rated as occurring simultaneously, the same rank numbers were replaced by the average rank. The ranks in each sequence were then normalized to the relative scaling of 0 to 1, so that “0” was assigned to the lowest rank and “1” to the highest rank (Germain & Laurin, 2009). Similarly, normalized ranks were calculated for ancestral developmental sequences reconstructed as described below.

### 2.4 Reconstruction of ancestral developmental sequences

To reconstruct ancestral developmental sequences, we used two different methods: event-pairing (Jeffery et al., 2002a) and continuous analysis (Germain & Laurin, 2009).

When event-pairing (Jeffery et al., 2002a) was used to infer ancestral sequences, all 171 pairs (19×18/2) of developmental events in each species were scored based on the relative timing; when one event occurred earlier, simultaneously, or later than another event, the timing was rated as 0, 1, or 2, respectively. By comparing the scored event-pairing matrices of different species, the ancestral event-pairing matrix was reconstructed at each node of the teleost phylogenetic tree in a parsimonious manner under accelerated transformation (acctran) and delayed transformation (deltran) optimizations using PAUP* software (Swofford, 2002). The ancestral event-pairing matrices were then reconverted to ancestral developmental sequences (Supplemental file 2).

In the continuous analysis (Germain & Laurin, 2009), the normalized ranks for each event calculated as described above were used as a continuous time scale to estimate the timing of the event in a hypothetical ancestor by squared-change parsimony (Maddison, 1991). The ranks of individual events in the hypothetical ancestors were estimated using the PDAP module of the Mesquite software (Maddison & Maddison, 2002, Midford, Garland & Maddison, 2003) (Figure S1-S19). According to the ranks, the events were then arranged into the ancestral developmental sequences (Supplemental file 2).

### 2.5 Quantification of rank variation for individual developmental events

To quantify rank variation for each event, the pairwise distance in the normalized ranks of an event was calculated by comparing the sequence on a branch to its immediate ancestor in the teleost phylogenic tree. The distances obtained from all the branches were then summed and averaged for the number of branches examined. The missing events in some branches were excluded from the calculation. The rates of evolution for developmental events by continuous analysis were obtained using the Mesquite PDAP module (Maddison & Maddison, 2002; Midford et al., 2003).

### 2.6 Detection of heterochronic shifts using a Parsimov algorithm

The heterochronic shifts between two developmental sequences at each phylogenetic node were estimated using the Parsimov algorithm developed by Jeffery et al. (2005). This parsimony-based algorithm determines the minimum number of event shifts that can explain the difference between two developmental sequences. Following the instructions, we implemented a Perl script, Parsimv7g.pl, with PAUP* output log file under both acctran and deltran optimizations, and mapped the detected heterochronic shifts onto the teleost phylogeny (Supplemental file 3).

### 2.7 Simulation of the reference distribution of heterochronic shifts

A reference distribution of heterochronic shifts over the teleost phylogeny was created by computer simulation based on a simple null assumption that a heterochronic shift occurs at a constant rate per unit time and accumulates in proportion to branch length. In this simulation, we did not consider the event-dependent differences in the shift frequencies. The simulation distributed the estimated number of heterochronic shifts randomly over the teleost phylogenic branches depending solely on their branch lengths. The simulation was replicated 100,000 times to obtain a reference null distribution. The null distribution of heterochronic shifts was then compared to the distribution of heterochronic shifts estimated from experimental datasets.

### 2.8 Calculation of total rank changes in individual phylogenic branches

The pairwise distance in the normalized ranks for each event was calculated as described above. The distances for all the events were then summed for each branch. The calculated values were then normalized by dividing by the number of examined events (excluding the missing event), and represented as the total rank changes per branch (Supplemental file 4).

## 3. Results

### 3.1 Phylogenetic relationship of the 31 actinopterygian species

For the present analyses, we selected 30 teleost species belonging to 13 distinct orders as the in-group because their developmental sequences have been well-documented (Figure 1). As an out-group, we used the amiadae fish, *Amia calva*, as a recent molecular genetic analysis had confirmed its location as the out-group of the teleost (Near et al., 2012); furthermore, it retains ancestral morphological characteristics such as mineralized scales, gas bladder lung, and heterocercal tail (Nelson, Grande & Wilson, 2016). In the constructed phylogenetic tree, the examined species were widely distributed and represented distinct branches of the teleost clade (Figure 2). Because there are very few reports on fish development in a marine environment, the species covered in this study were basically freshwater teleosts; however, several anadromous species were also included such as *Gasterosteus aculeatus*, which develops in freshwater but alternates between the sea and freshwater as an adult.

**FIGURE 2.**
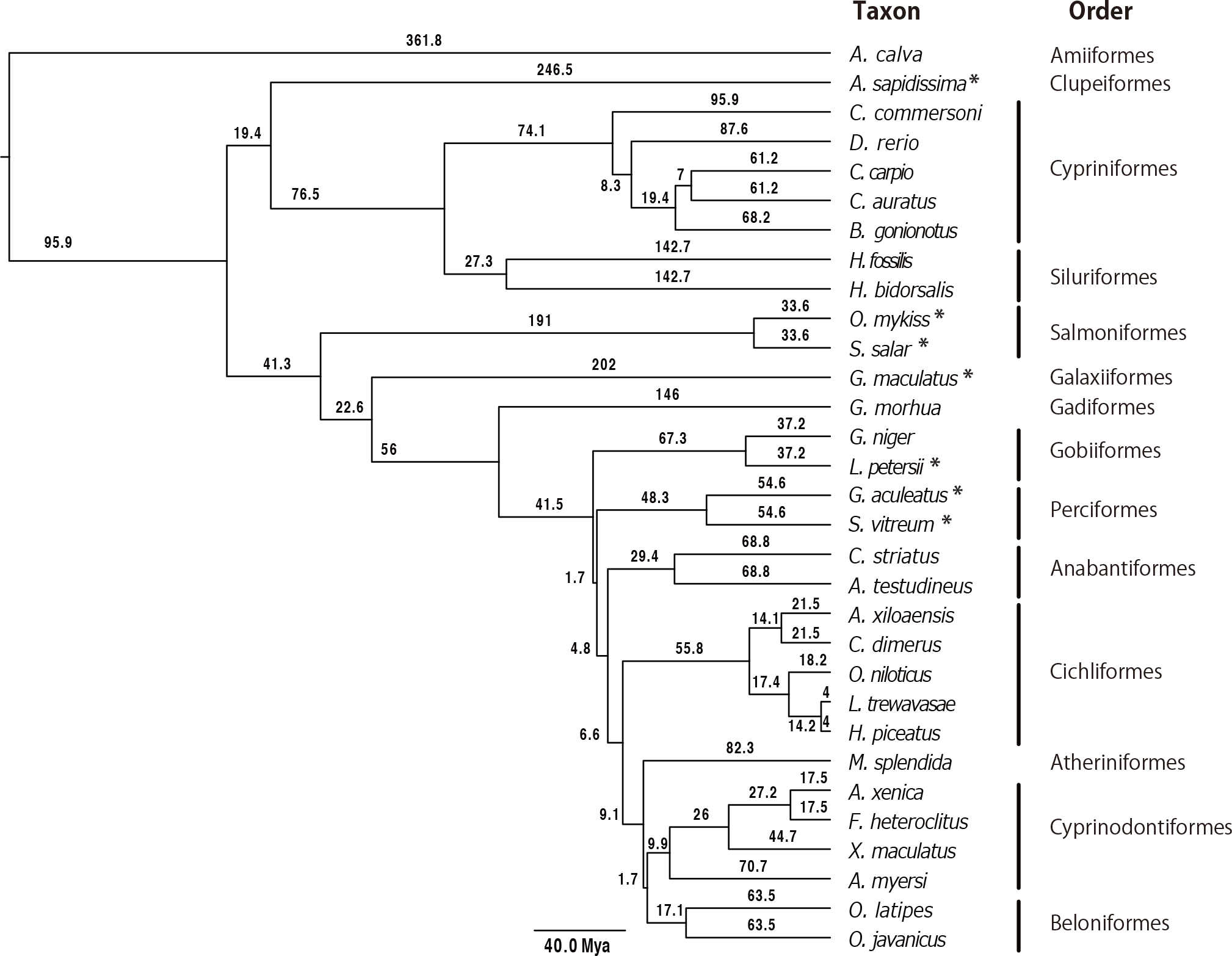
Phylogenetic relationships of the 31 actinopterygian species. The phylogenetic tree of the 31 actinopterygian species examined in this study. The asterisk marks anadromous teleosts, while all the others are freshwater teleosts. The numbers beside the branches indicate the divergent times (Mya). The information sources for tree construction are as follows: Near et al. (2012, 2013), Saitoh et al. (2011), Yang et al. (2015), Perez et al. (2007), Friedman et al. (2013), Pohl, Milvertz, Meyer & Vences (2015).

### 3.2 Comparison of temporal orders of developmental events among teleost species

We selected 19 developmental events that appeared consistently as landmarks in the developmental staging of many teleost species (Figure 1). To gain an overall view of developmental sequences involved in the formation of entire body, we intentionally included events that belong to substantially different biological systems and contexts such as those originating from different germ layers or giving rise to different cell types or separate body parts. Additionally, the list also included a small number of interrelated events such as formation of the optic vesicles/placodes/primordia (op), lenses or lens placodes (le), and eye pigmentation (ep). We gathered information on these 19 events from articles reporting the development of 31 actinopterygian species, and ranked the orders of individual events in the temporal sequence for each species (Figure 1).

We first compared rank orders of individual events among 30 in-group teleost species. To minimize effects of simultaneous occurrence of events and missing data, the raw ranks (Figure 1) were rescaled to the normalized ranks that fit within the same range, from 0 to 1, in all the teleost species (see Methods). Figure 3a shows the distribution of the normalized ranks for individual developmental events, which were horizontally arranged according to the average values. Interestingly, the ranges of variation in the ranks differed widely, depending on the event. One extreme case was embryonic shield (es), which always appeared first and with no variation in the developmental sequences obtained from the 29 teleost species, excluding one missing description for *Galaxias maculatus* (Figure 1). In contrast, relatively large variations in the rank were observed for the appearance of olfactory vesicles/pits/primordia (olf) and medial finfold (mff), suggesting that the temporal order of these events in the developmental sequence can change more easily (Figure 3a).

**FIGURE 3.**
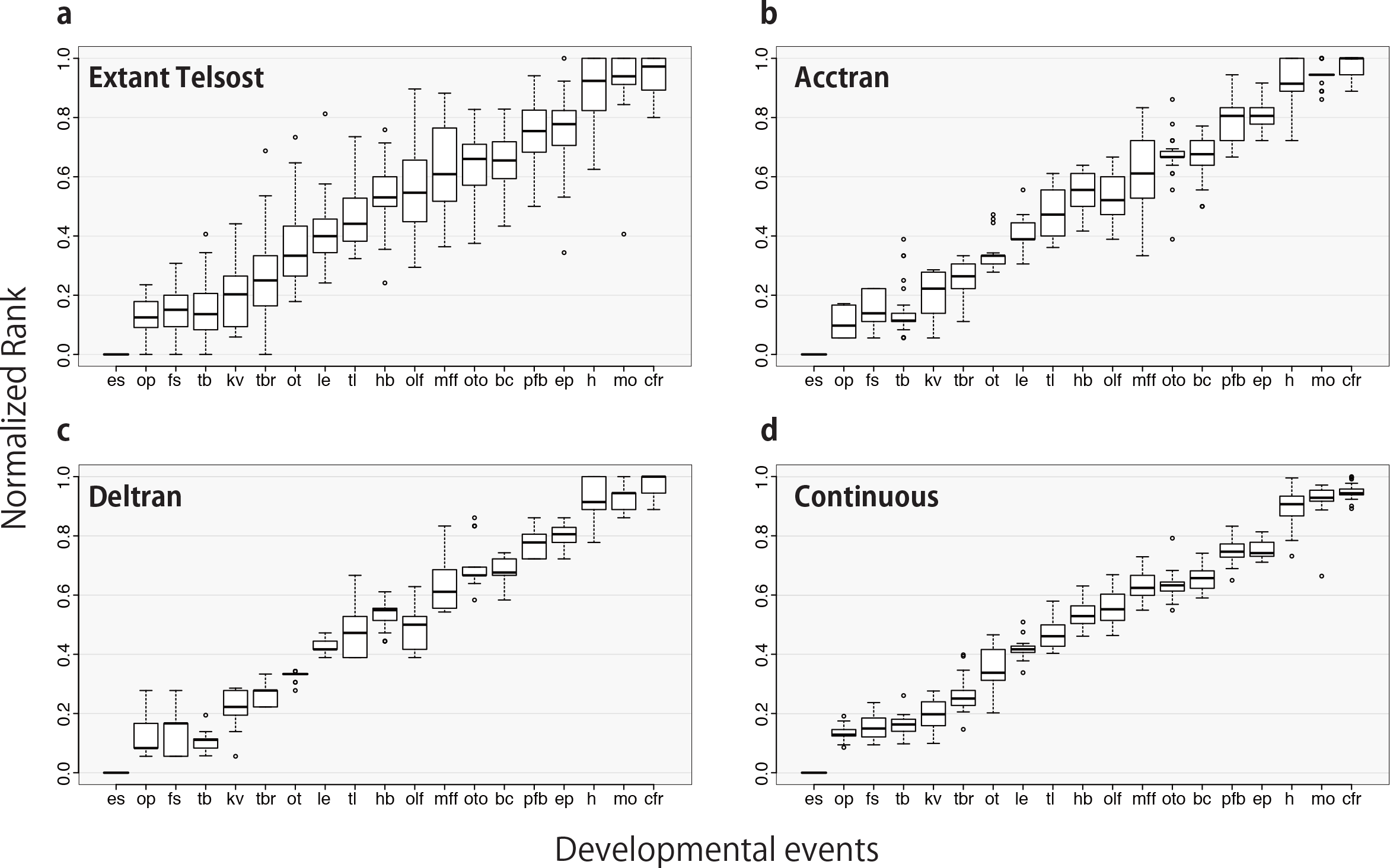
Distribution of ranks of events in the developmental sequence. The boxplot shows the statistical distribution (minimum, first quartile, median, third quartile, maximum, and outliers) of normalized ranks for individual developmental events obtained from the extant 30 teleost in-group data (a), and ancestral developmental sequences reconstructed by acctran (b) and deltran (c) optimizations and continuous analysis (d). In all the panels, the developmental events are aligned horizontally from left to right in the same order according to the average ranks in the extant fish sequences.

To explore the evolutionary history of developmental sequences, we next estimated ancestral developmental sequences on each phylogenetic branch using the event-pairing method (Jeffery et al., 2002a). This algorithm compares the relative orders of all the event pairs between two different developmental sequences and generates the ancestral sequences determined as a parsimonious solution under acctran and deltran optimizations (Supplemental file 2). Using the obtained ancestral developmental sequences, we compared the normalized ranks of individual events as in Figure 3a. Overall, the rank orders of individual events in the ancestral developmental sequences (Figure 3b and 3c) were similar to the extant sequences (Figure 3a). When the developmental events were horizontally aligned in the same order as the extant average ranks, there were only a few inversions in the order of two successive events at the average level (e.g., the order between first somite [fs] and tail bud [tb]). The range of variation for individual events was also similar to the extant sequences, further confirming that the ranks of some developmental events change more frequently than others during evolution.

We repeated the same rank analyses using ancestral developmental sequences inferred by a different method developed by Germain & Laurin (2009). This “continuous analysis” uses the normalized rank of events as a continuous ontogenetic time scale and directly estimates the timing of individual events that should occur in hypothetical ancestors based on squared-change parsimony (Maddison, 1991). Because the two methods are based on different assumptions, they produced significantly different ancestral sequences (Supplemental file 2). The rank variations of the sequences showed a similar but slightly different trend, namely that the ranks for appearance of otic vesicles/placodes/primordia (ot) and olfactory vesicles/pits/primordia (olf) were relatively highly variable (Figure 3d).

### 3.3 Evaluation of rank changes through the evolutionary history of the teleost

Because rank variations fluctuate depending on the event, we next evaluated rank changeability phylogenetically along the course of teleost evolution. We measured the pairwise rank distance between a sequence and its immediate ancestral sequence across the node in the teleost phylogenic tree (Supplemental file 4), represented as the average value per branch (Figure 4a). For this analysis, ancestral developmental sequences were obtained by event-pairing (Jeffery et al., 2002a) under acctran and deltran optimizations. The index scores were highly consistent regardless of the optimization method used (Spearman’s rank correlation for the two optimizations; r = 0.7621). When the events were chronologically arranged along the standard ontogenic time frame, defined as the average rank orders in the extant teleosts, the overall shape of the ontogenic histogram did not show any clear characteristics in the middle phase of the developmental sequence characterized by three brain regionalization (tbr), otic vesicles/placodes/primordia (ot), and lenses or lens placodes (le). This developmental term typically corresponds to the narrow neck in the hourglass model. The ontogenic histogram was instead relatively skewed toward the late tail bud stage involving olfactory vesicles/pits/primordia (olf) and otoliths (oto). Because the late tail bud stage also included low-profile events such as blood circulation (bc), we do not consider the weak upward trend as a continual ontogenic pattern formed by temporally clustered events.

**FIGURE 4.**
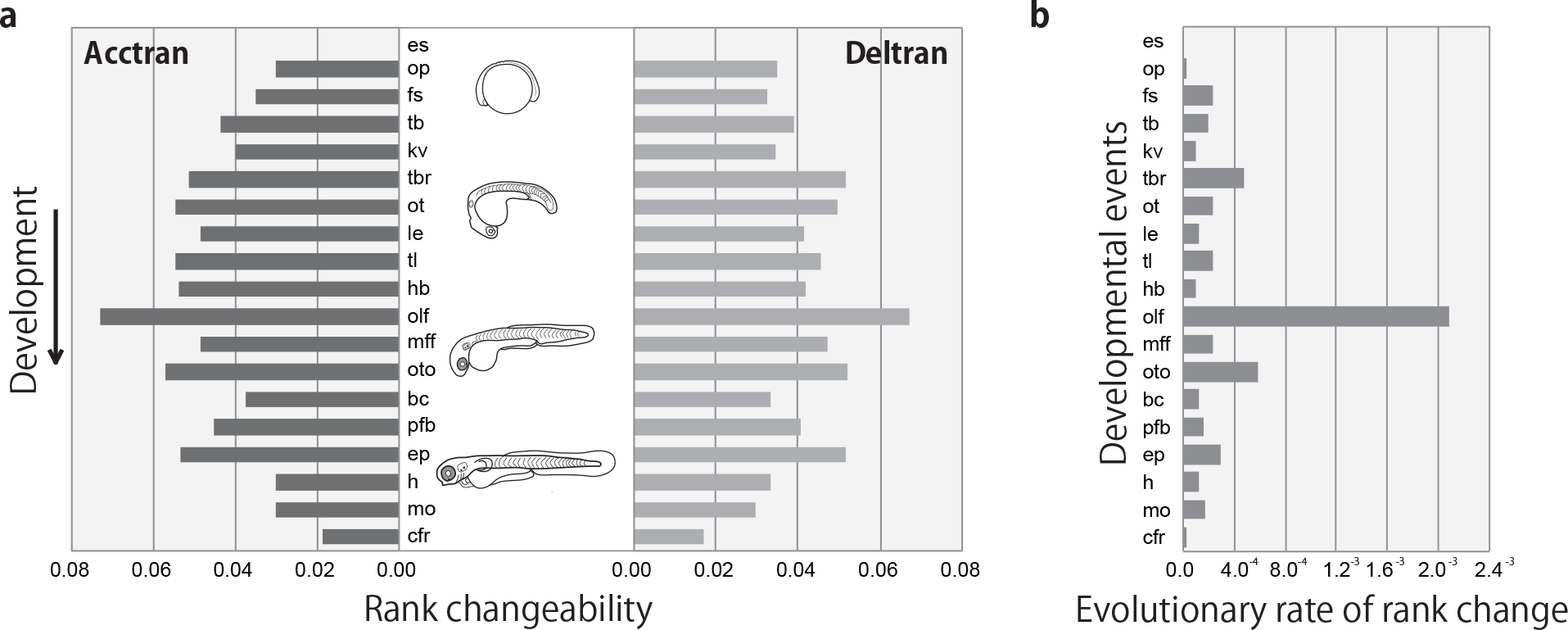
Rank changeability of individual developmental events during evolution. (a) Rank variation is shown as the average of pairwise rank distances calculated by comparing the pair of a sequence and its immediate ancestral sequence. The ancestral developmental sequences were estimated using acctran (left) and deltran (right) optimizations of the event-pairing method (Jeffery et al., 2002a). The events are vertically arranged from top to bottom along the standard ontogenic time frame defined by the average developmental sequence in the extant teleosts (Figure 2). (b) The evolutionary rate of rank changes for individual events obtained by continuous analysis (Germain & Laurin, 2009). The events are arranged in the same order as in (a).

We also examined evolutionary rank changes using a different index. Based on continuous analysis (Germain & Laurin, 2009), the evolutionary rate of rank changes can be obtained for each event (Figure 4b). This index shows rank changeability per million years and therefore differs slightly from the one per branch used in the previous analysis. The calculation also involved the set of different ancestral sequences estimated by continuous analysis. Nevertheless, this new index was basically similar to the previous analysis, with the olfactory vesicles/pits/primordia (olf) exhibiting high rank changeability. In the ontogenic sequence, this high score event was like an independent peak with no indication of a global ontogenic pattern. These and previous results further suggest that the ranks of olfactory vesicles/pits/primordia (olf) have changed more frequently during evolution than the other events.

### 3.4 Event exchange rates in developmental sequences

We next focused on the actual sequence order of developmental events. Figure 5 shows the percentage of sequences in which one event (shown in the row) occurs later than another (shown in the column) among the 30 extant in-group species. In general, the sequence of two temporally distant events was somewhat conservative, with no order reversal with many combinations, whereas the orders of neighboring events changed more frequently. Closer inspection of anatomically interrelated events indicates that the temporal order of the optic vesicles/placodes/primordia (op) and the lenses or lens placodes (le) was fixed in all the species, and that of the lenses or lens placodes (le) and the eye pigmentation (ep) was practically fixed, except for one sequence reversal in *Heterobranchus bidorsalis*. Similar results were obtained from the comparison of event orders in the ancestral developmental sequences (Figure S20a and S20b).

**FIGURE 5.**
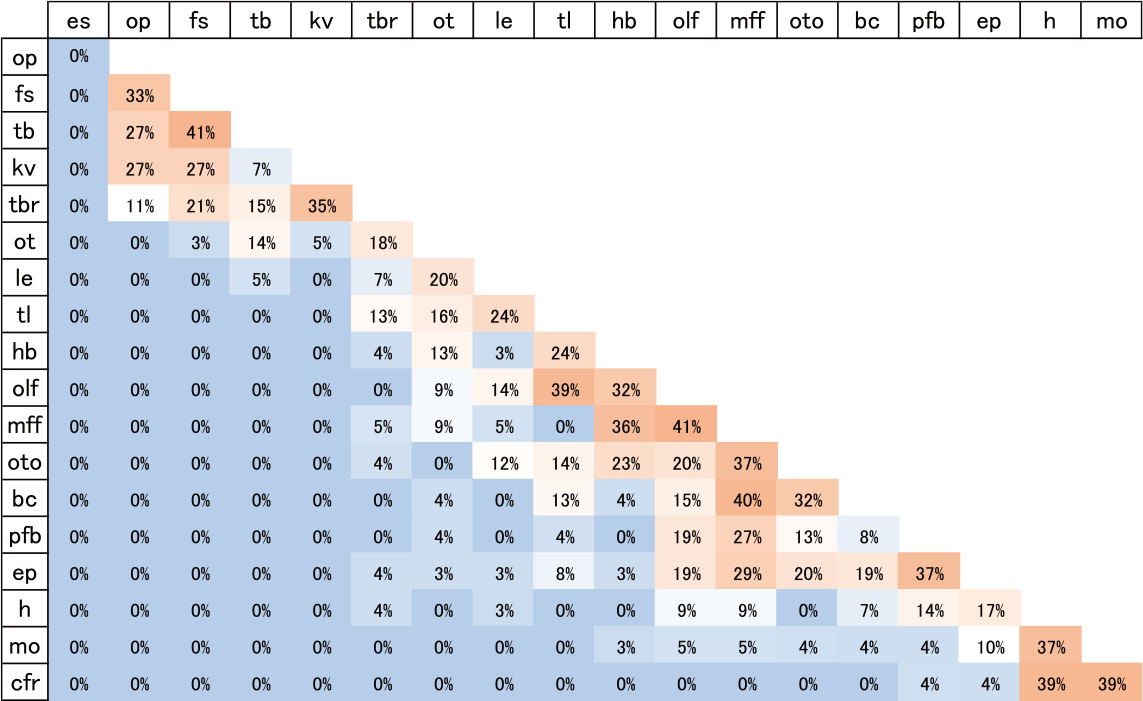
Sequence orders of event pairs in extant developmental sequences. The event sequence matrix represents all the pairwise combinations of developmental events. The number shows the percentage of the sequences in which the row event occurred later than the column event, and was calculated from the dataset of 30 extant teleost species, excluding the missing event data. The individual cells are heatmap color-coded according to percentage value.

### 3.5 Distribution of heterochronic shifts across the teleost phylogenetic tree

Using event-pairing with a Parsimov algorithm (Jeffery et al., 2005), we next searched for heterochronic shifts of the events that can explain the changes from one sequence to another at every node of the teleost phylogenetic tree. Although this is a parsimony-based algorithm and, therefore, estimates the minimum number of event shifts, we obtained 167 (acctran), 162 (deltran), and 91 (conserved between acctran and deltran) heterochronic shifts in total (Supplemental file 3). The teleost phylogenetic tree has 59 branches in total and when the estimated shifts were mapped onto the tree, multiple heterochronic shifts were observed in almost all the branches (Supplemental file 3).

Because a substantial number of heterochronic shifts were inferred, we examined the distribution of these shifts in the teleost phylogeny. Based on the simple null hypothesis that a heterochronic shift occurs at a stochastically random manner and accumulates neutrally and exponentially along phylogenic time, we calculated a reference distribution of heterochronic shifts by computer simulation (white circles in Figure 6a and 6b). Compared to this reference null distribution, the actual heterochronic shifts inferred by the Parsimov algorithm exhibited a somewhat enigmatic distribution, with a constant number of shifts relative to branch length with both acctran and deltran optimizations (black circles in Figure 6a and 6b). The number of heterochronic shifts per branch showed a smaller coefficient of variation compared to the reference value (Figure 6c), indicating that branch-by-branch fluctuations in the number of heterochronic shifts were more limited than those estimated from random allocation of the shifts. The constant trend in the shift number was also supported by the fact that the number of phylogenetic branches harboring no heterochronic shifts from the experimental dataset was significantly smaller than the reference data (Figure 6d). Because inclusion of an extremely long branch could skew the statistical results, we performed the same statistical comparison but using only relatively short branches (≤ 50 Mya and ≤20 Mya in Figure 6c, 6d). We also replaced the ≤ phylogenic time scale with the generation number (Figures S21 and S22) by considering the average generation time of individual species (Supplemental file 5). However, these modifications did not qualitatively change the results of the analysis.

**FIGURE 6.**
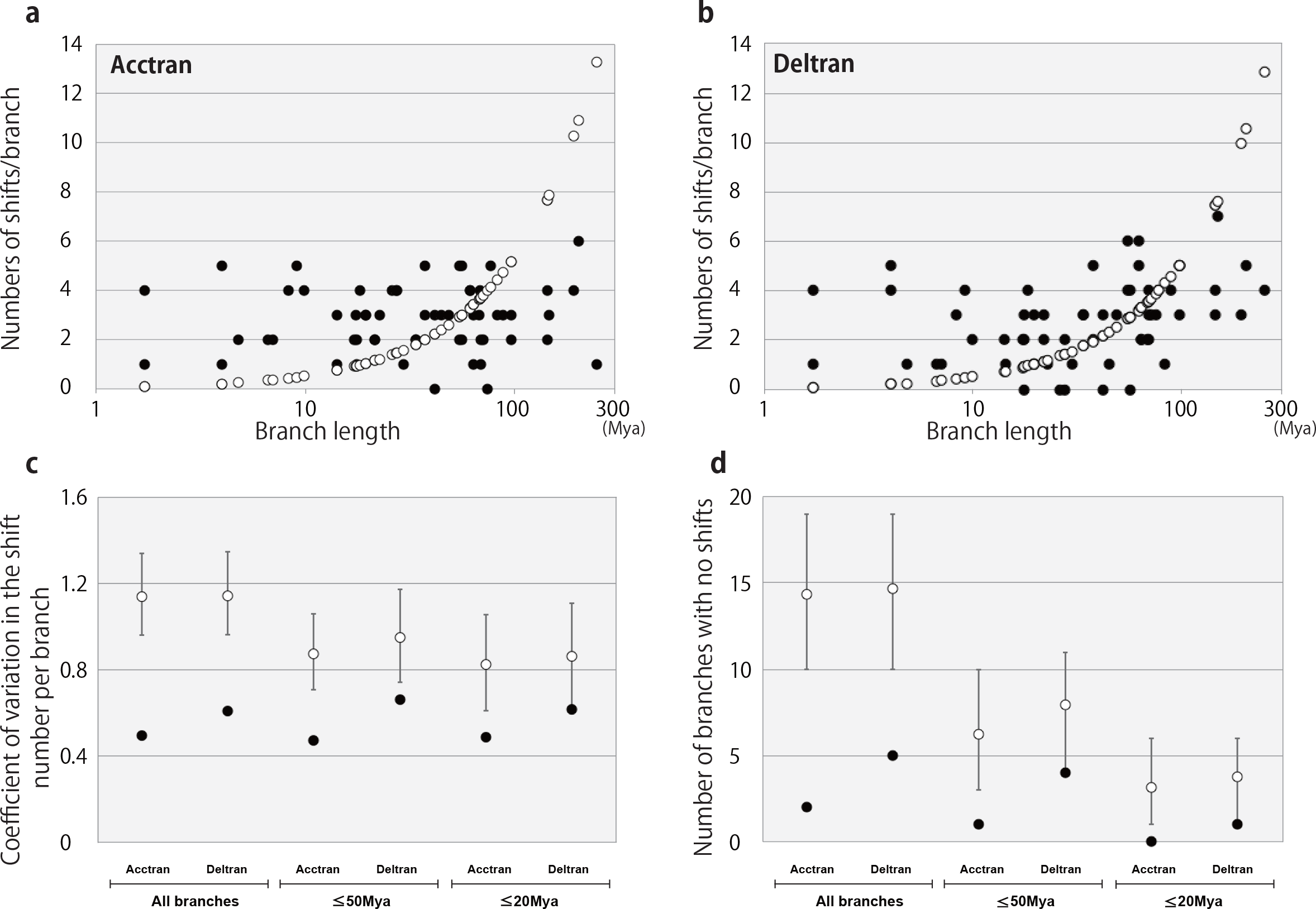
Distribution of heterochronic shifts estimated by event-pairing in the teleost phylogeny. (a,b) Scatter plots showing the relationship between the phylogenetic branch length and number of heterochronic shifts estimated from the extant and ancestral developmental sequences by event-pairing with acctran (a) and deltran (b) optimizations (black circle). The reference distribution (open circles) hypothesizes a linear correlation between the branch length and shift number. (c) The coefficient of variation for the number of heterochronic shifts in a branch. The black and open circles show the observed and reference values, respectively. The vertical bars three different groups: all branches, branches shorter than 50 million years (Mya), and branches shorter than 20 Mya. (d) The number of branches without heterochronic shifts calculated from observed (black circle) and reference (open circle) data in three different branch length categories, as explained in (c). Vertical bars indicate 95% confident intervals of the reference value. The number of branches in each analysis were 59 (all branches), 33 (shorter than 50 Mya), and 19 (shorter than 20 Mya).

A potential caveat about standard parsimony is that it may give ambiguous solutions as equally likely by considering only the number of changes, and not the branch length. Thus, we conducted a similar analysis of heterochronic rank changes using the ancestral developmental sequences inferred by continuous analysis, a modified parsimony method that incorporates branch length information to find a single most likely solution (Germain & Laurin, 2009). When we used the ancestral sequences inferred by standard parsimony-based event-pairing, the calculated total rank changes involving all the events per branch exhibited a constant trend against the branch length (Figure 7a and 7b, Supplemental file 4), similar to the preceding analysis. The Spearman’s rank correlation coefficients with regard to branch length showed a marginally significant, but weak correlation (acctran: 0.3040, deltran: 0.3428). When the ancestral sequences inferred by continuous analysis were used instead for the calculation, the total rank changes presented a stronger correlation with the branch length, showing a clear proportional increase (Figure 7c, Spearman’s rank correlation coefficient: 0.5723). These results imply that the selection of ancestral sequences has a strong influence in this type of analysis, and that the enigmatic distribution of heterochronic event shifts observed with standard parsimony can be corrected by allowing for the effect of phylogenetic time, leaving the distribution of event shifts still unclear. A more important indication from the comparisons may be that similar numbers of heterochronic event changes were estimated, regardless of the method (Figure 7d and 7e). These numbers, which are parsimonious estimates, further support that heterochronic shifts of developmental events were frequent during teleost evolution.

**FIGURE 7.**
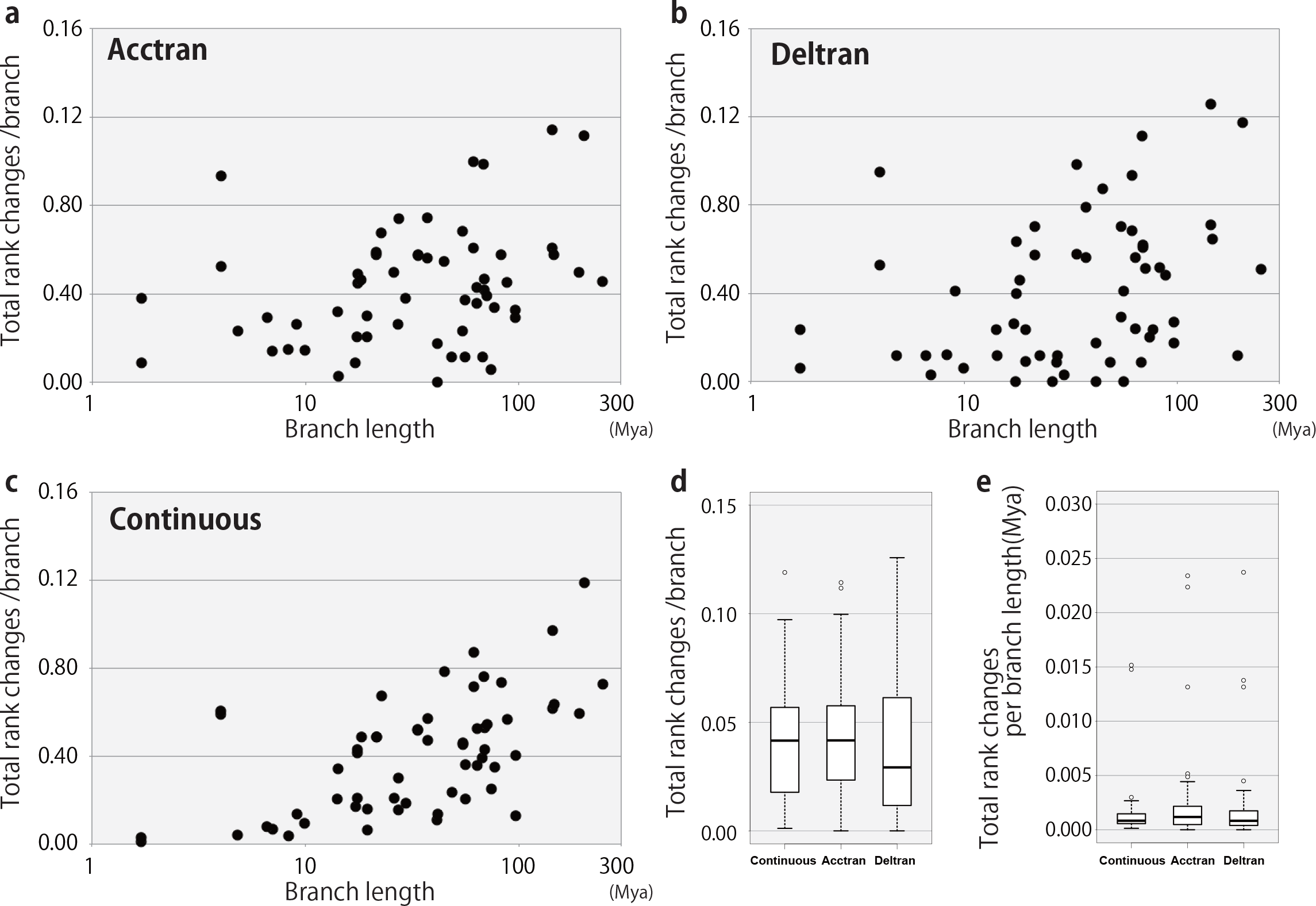
Comparison of the distribution of heterochronic rank changes estimated by different methods. (a to c) Scatter plots showing the relationship between branch length and the amount of total rank changes estimated to have occurred in the branch. The ancestral developmental sequences were estimated by event-pairing with acctran (a) and deltran (b) optimization, and continuous analysis (c). (d) The number of total rank changes divided by the number of branches. The coefficients of variation: 0.6405 (continuous), 0.6130 (acctran), and 0.8116 (deltran). (e) The number of total rank changes per branch per million years. The coefficients of variation: 1.7396 (continuous), 1.8769 (acctran), and 1.9670 (deltran).

## 4. Discussion

In the present study, we systematically analyzed evolutionary changes in the developmental sequences of teleosts using a wide range of species and developmental events. The results revealed how teleost ancestors experienced dynamic and frequent rearrangements of developmental sequences leading up to extant species. This first comprehensive analysis of teleost developmental sequences will provide a valuable foundation toward understanding the significance of evolutionary changes in developmental sequences for animal morphological diversification.

Initially, we expected a more conservative evolution of developmental sequences because we only examined highly conserved traits in the phylogeny and species belonging to a restricted infraclass. However, we found more frequent heterochronic changes than expected, leading to very weak phylogenetic signals. This may be due to polymorphic fluctuations in developmental timing, observable even in a single species (de Jong, Colbert, Witte & Richardson, 2009; Mitgutsch, Wimmer, Sanchez-Villagra, Hahnloser & Schneider, 2011). If the developmental timing is not fixed, comparisons of different species staged under different conditions should further increase false positive signals (Weisbecker & Mitgutsch, 2010). In addition, our wide selection of developmental events may have influenced the results; in this study, we intentionally adopted events covering a variety of embryonic origins, cell types, body parts, or biological systems to understand global body patterning. In contrast, several previous studies aimed for in-depth understanding the developmental sequences for a specific body part or organ (Schlosser, 2008; Hautier et al., 2011). For example, in the comparative analysis of neural development in 18 mammalian species, the whole sequence of 271 neural developmental events was strikingly preserved, even though many of the events occur in separate places (Workman, Charvet, Clancy, Darlington & Finlay, 2013). The ossification sequence of cranial bones in early mammals was also seen to be highly conserved (Sanchez-Villagra et al., 2008). It is conceivable that development of individual elements in the same tissue or organ is more hierarchically constrained, whereas different tissues or body parts may have more freedom to behave as independent modules during development (Klingenberg, Badyaev, Sowry & Beckwith 2001; Schmidt & Starck, 2010; Kawanishi et al., 2013; Laurin, 2014). The modular nature of individual body parts may explain why relatively frequent changes of developmental sequences were observed in this study, and may possibly serve as a source for the independent evolution of different body parts.

Another important note from the present study is that heterochronic shifts were frequent but not random. The rank analyses indicated that the temporal orders of some developmental events change more drastically than others during evolution. Although the magnitude fluctuates according to the method, the olfactory vesicles/pits/primordia (olf) consistently exhibited high rank changeability in all the analyses. The olfaction is a critical sensor for environment monitoring in animals. Conceivably, ancestral teleosts could have benefited from temporally accelerating or decelerating the formation of the olfactory sensory organs for better adaptation to environmental conditions.

Regarding the phylotypic period, the present study did detect neither a significant signature of evolutionary restriction of ontogenic event changes, nor an inverted increasing trend of heterochronic shifts during the middle phase of development, as previously reported (Bininda-Emond et al., 2003). Overall, we did not observe ontogenically clustered behavior of developmental events. Conceptually, our finding is close to the conclusion of Poe & Wake (2004): the simple rule that the orders of temporally neighboring events change more frequently is more relevant than other complicated models. Our result is important because we used different datasets for this analysis.

Although maximum parsimony has been a widely used criterion in evolutionary studies, it contains various drawbacks (Germain & Laurin, 2009). One drawback in the present study was that it considers only the number of character changes, and not phylogenetic time, to calculate the ancestral sequence. For example, under maximum parsimony, a single change on one short internal branch is always chosen as the realistic scenario compared to two parallel changes on two long external branches (Felsenstein, 1978). We obtained a somewhat enigmatic distribution of heterochronic shifts using the sequences inferred by the standard parsimony-based event-pairing; however, a more reasonable distribution was obtained using sequences inferred by continuous analysis, a method that is also parsimony-based but also incorporates branch length estimates. Thus, in our analyses, the latter method may be more appropriate for estimating the ancestral sequences. The results also indicated that the potential distribution of heterochronic shifts over the phylogenic tree is arbitrary and depends on the optimization protocol. To assess the evolutionary distribution of heterochronic shifts, a more effective high-resolution approach is needed. A potential alternative would be a simulation-based approach combining a stochastic model with parsimony calculation. This would require rigorous calculations and considerations but deserve a future challenge.

It is not clear whether teleost-specific circumstances are reflected in the present results. For example, external temperature is known to affect developmental time frames (Mabee, Olmstead & Cubbage, 2000; Schmidt & Starck, 2010) and, as most teleosts reproduce by external fertilization and the embryos develop under fluctuating temperatures, temporal shifts in individual developmental events may sometimes occur in teleosts in a natural environment. Accordingly, teleosts may be naturally accustomed to sporadic shifts in developmental events that may increase the probability of a shift being adopted in a persistent manner. Future analyses using other groups of animals will address the specificity and generality of the current results in evolutionary development.

## Acknowledgements

We thank Dr. Toshihiko Shiroishi, Dr. Yasushi Hiromi, Dr. Naoki Irie, Dr. Takanori Amano, Dr. Yuuta Moriyama, and Dr. Kousuke Mouri for comments that helped to greatly improve the manuscript. We also thank Dr. Erin E. Maxwell, Dr. Luke B. Harrison, and Dr. Atsushi Kawakita for their advice on parsimony analyses. T.H. acknowledges the funding support from MEXT/JSPS KAKENHI Grants (17H05776, 17H05587, 16H04659). This work was supported in part by SOKENDAI (The Graduate University for Advanced Studies).

## Author contribution

F.I. designed the study, performed the majority of the experiments, analyzed data and wrote the manuscript. T.M contributed to designing and conducting the simulation. T.M. and T.H. supervised the study and helped F.I. to write the manuscript with input from other people.

