## Supplemental Figures for "Frequent Non-random Shifts in the Temporal Sequence of Developmental Landmark Events during Teleost Evolutionary Diversification"

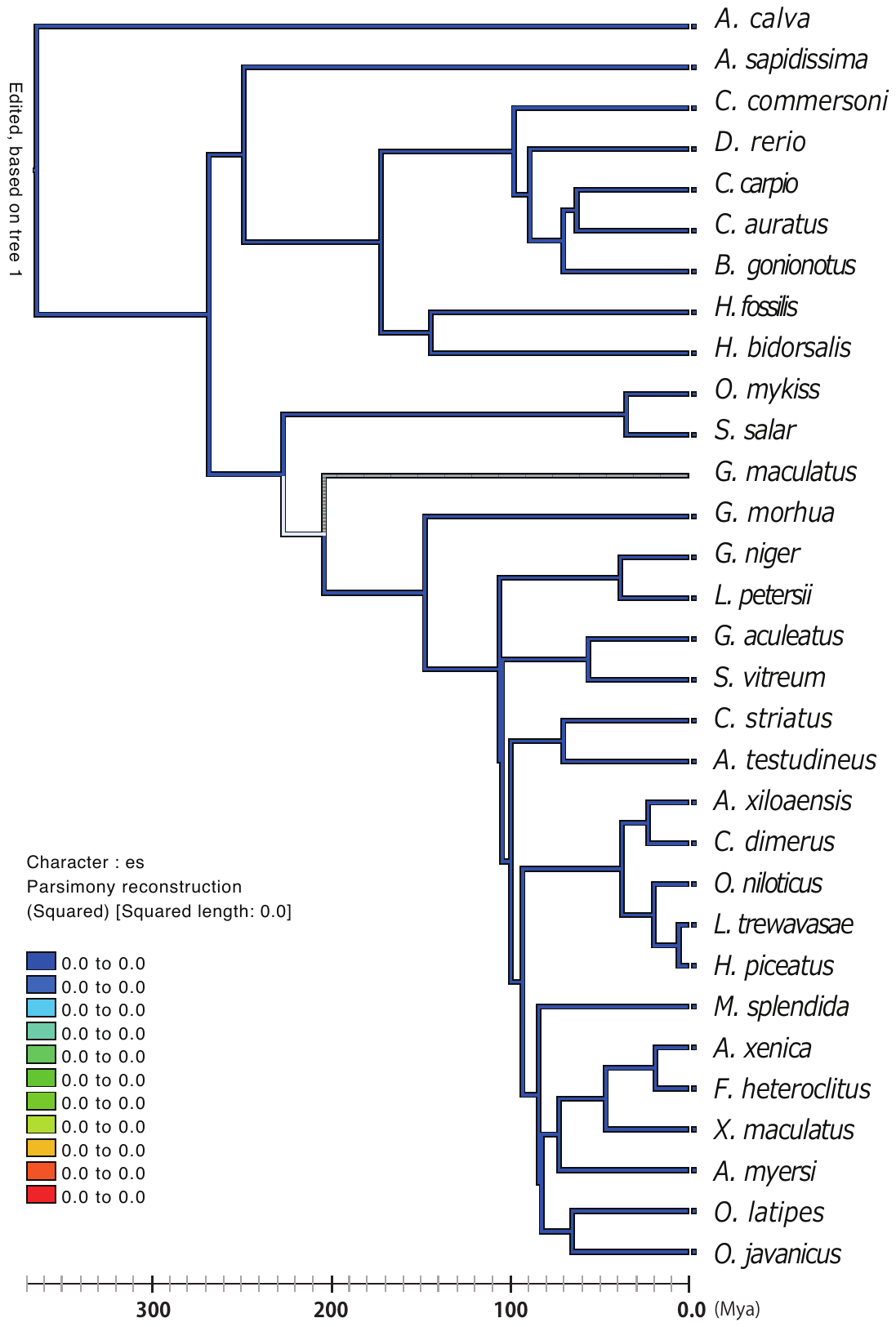

**Figure S1 Reconstructed ancestral ranks of embryonic shield (es) by continuous analysis**

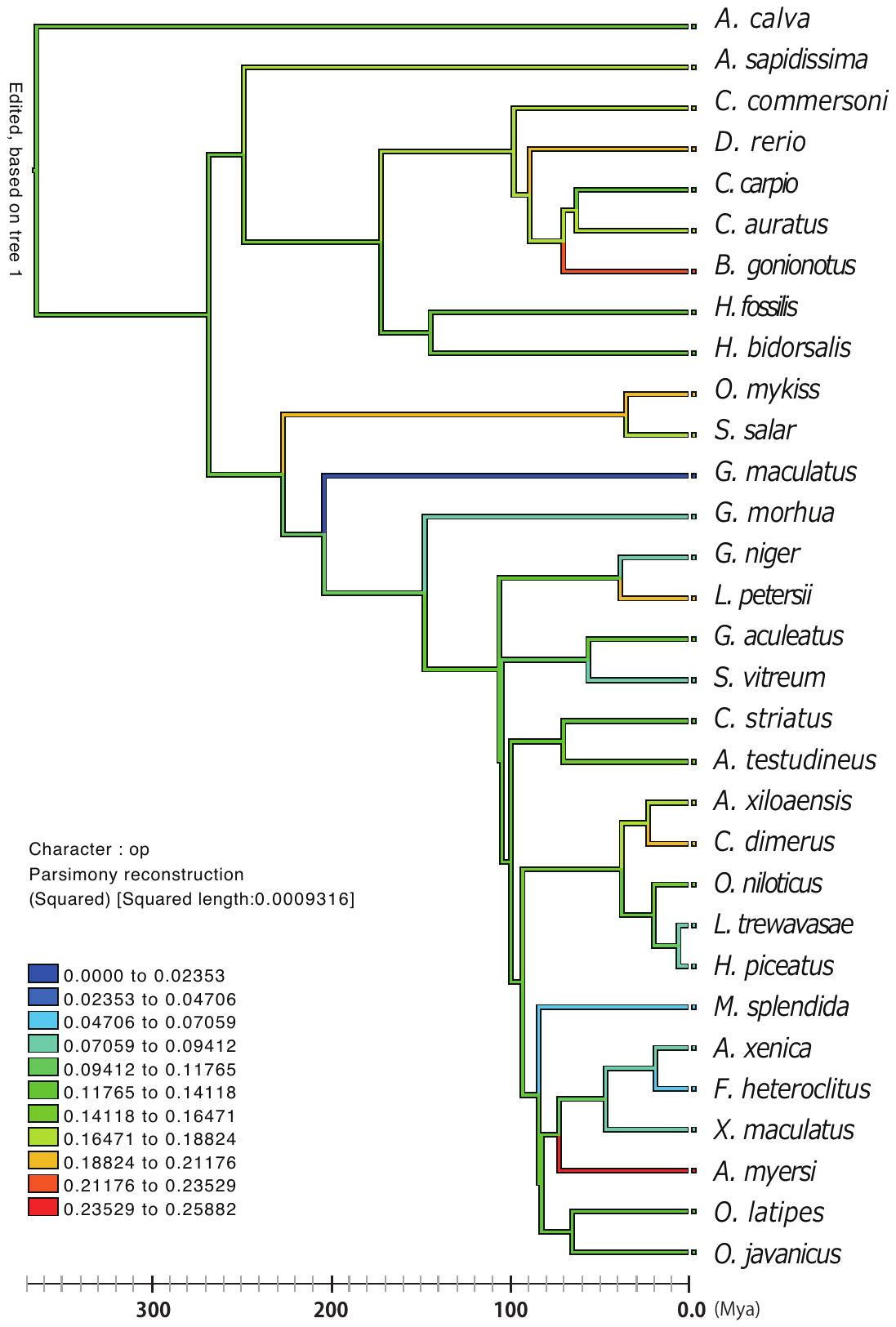

**Figure S2 Reconstructed ancestral ranks of optic vesicles/placodes/primordia (op) by continuous analysis**

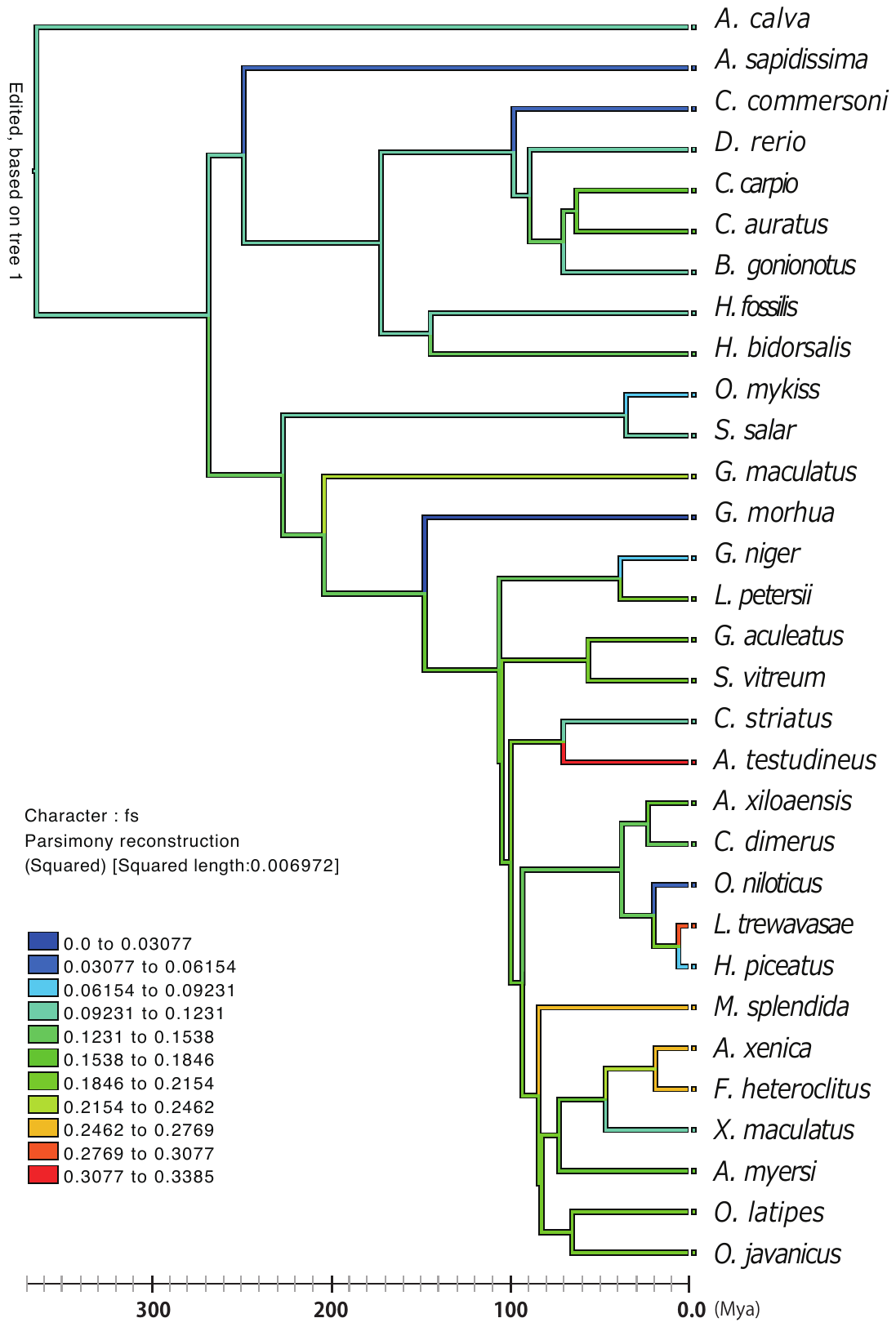

**Figure S3 Reconstructed ancestral ranks of first somite (fs) by continuous analysis**

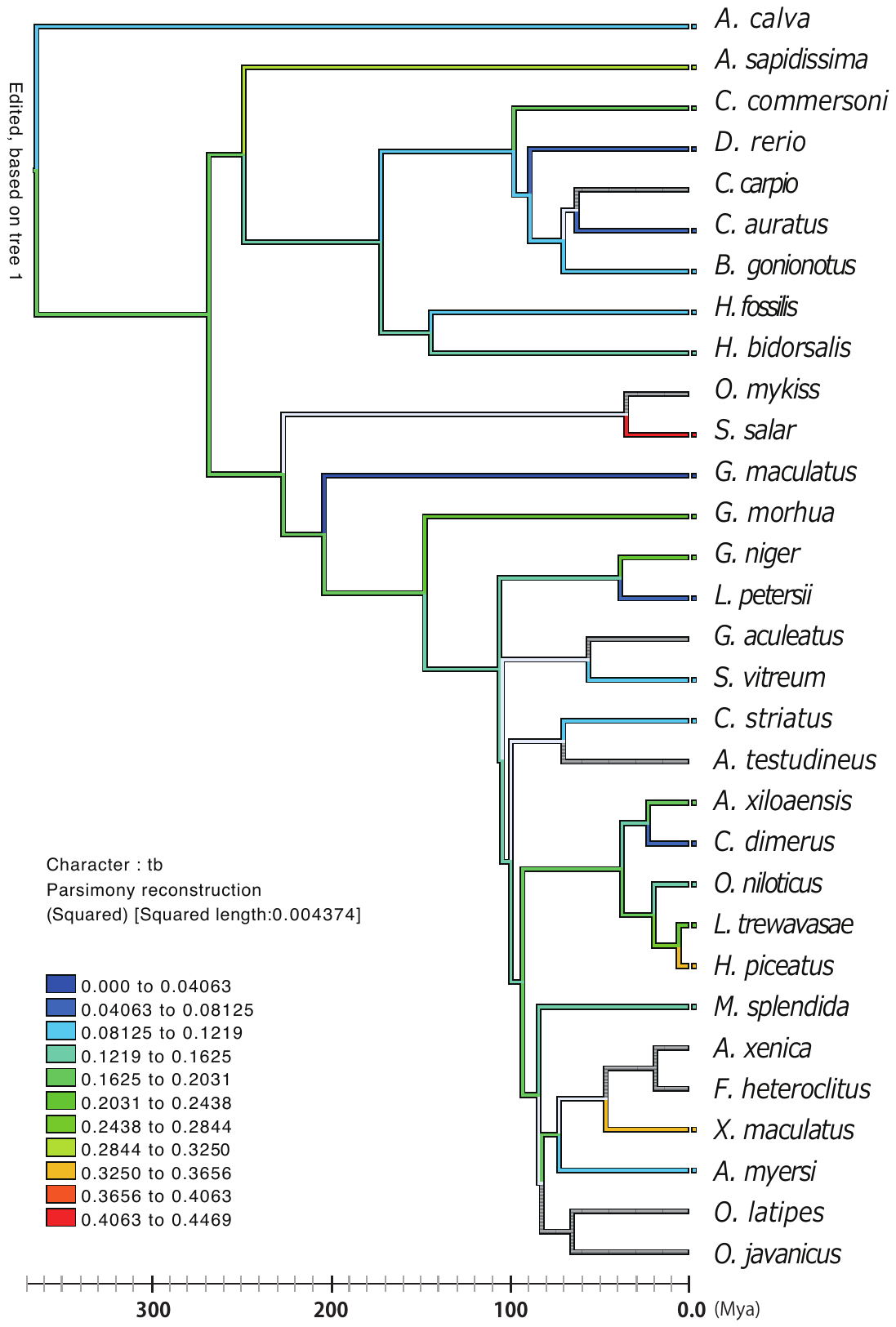

**Figure S4 Reconstructed ancestral ranks of tail bud (tb) by continuous analysis**

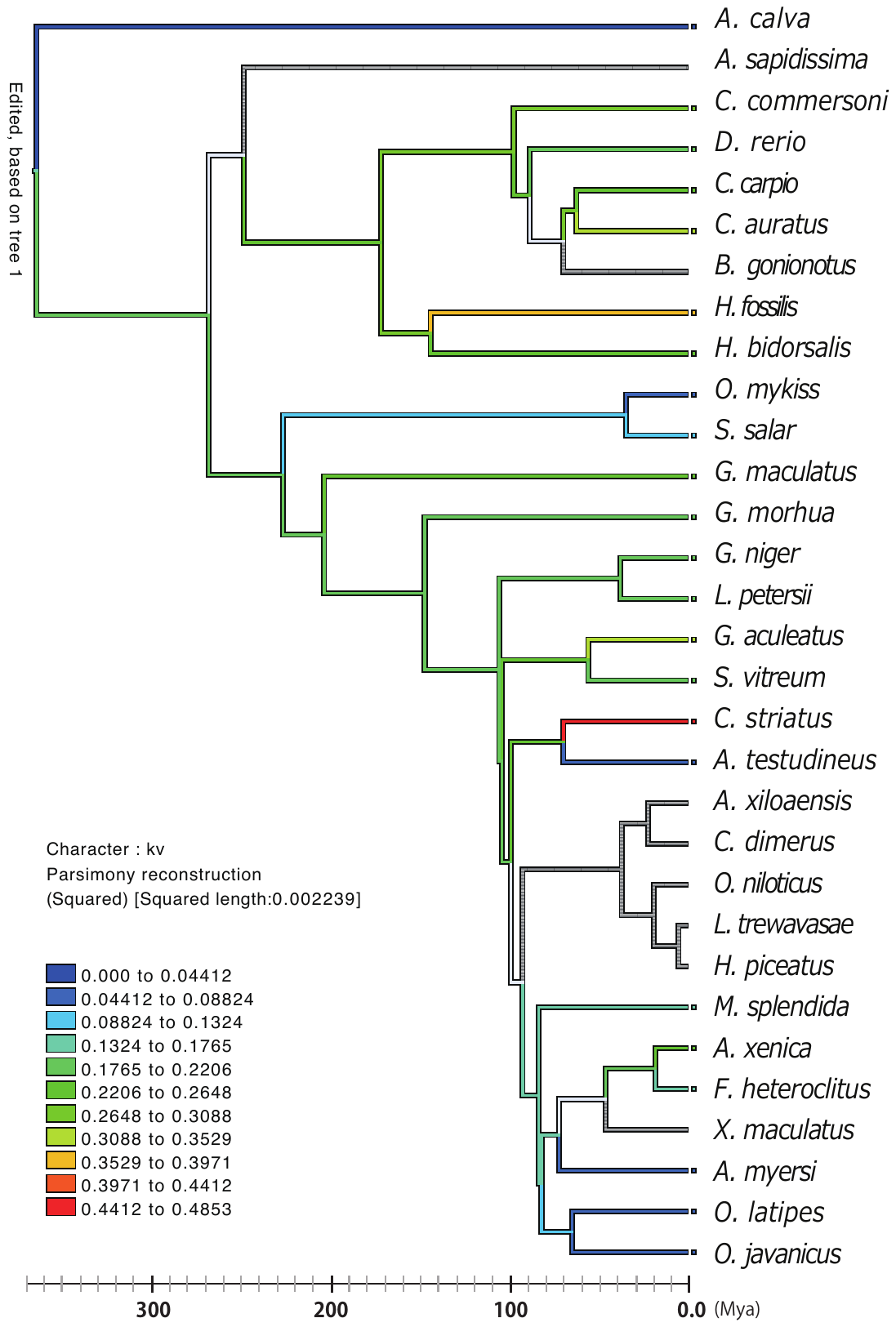

**Figure S5 Reconstructed ancestral ranks of Kupffer’s vesicle (kv) by continuous analysis**

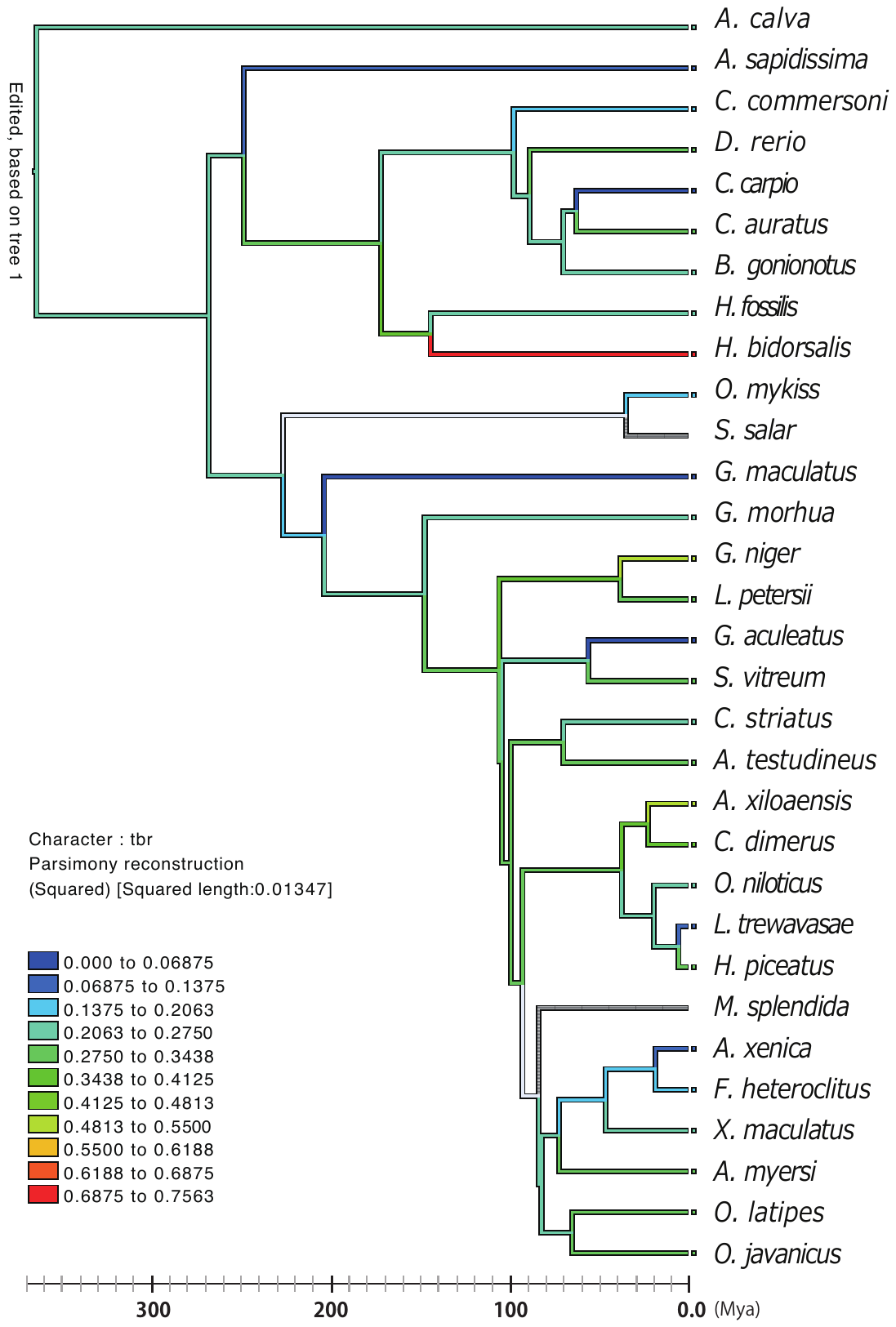

**Figure S6 Reconstructed ancestral ranks of three brain regionalization (tbr) by continuous analysis**

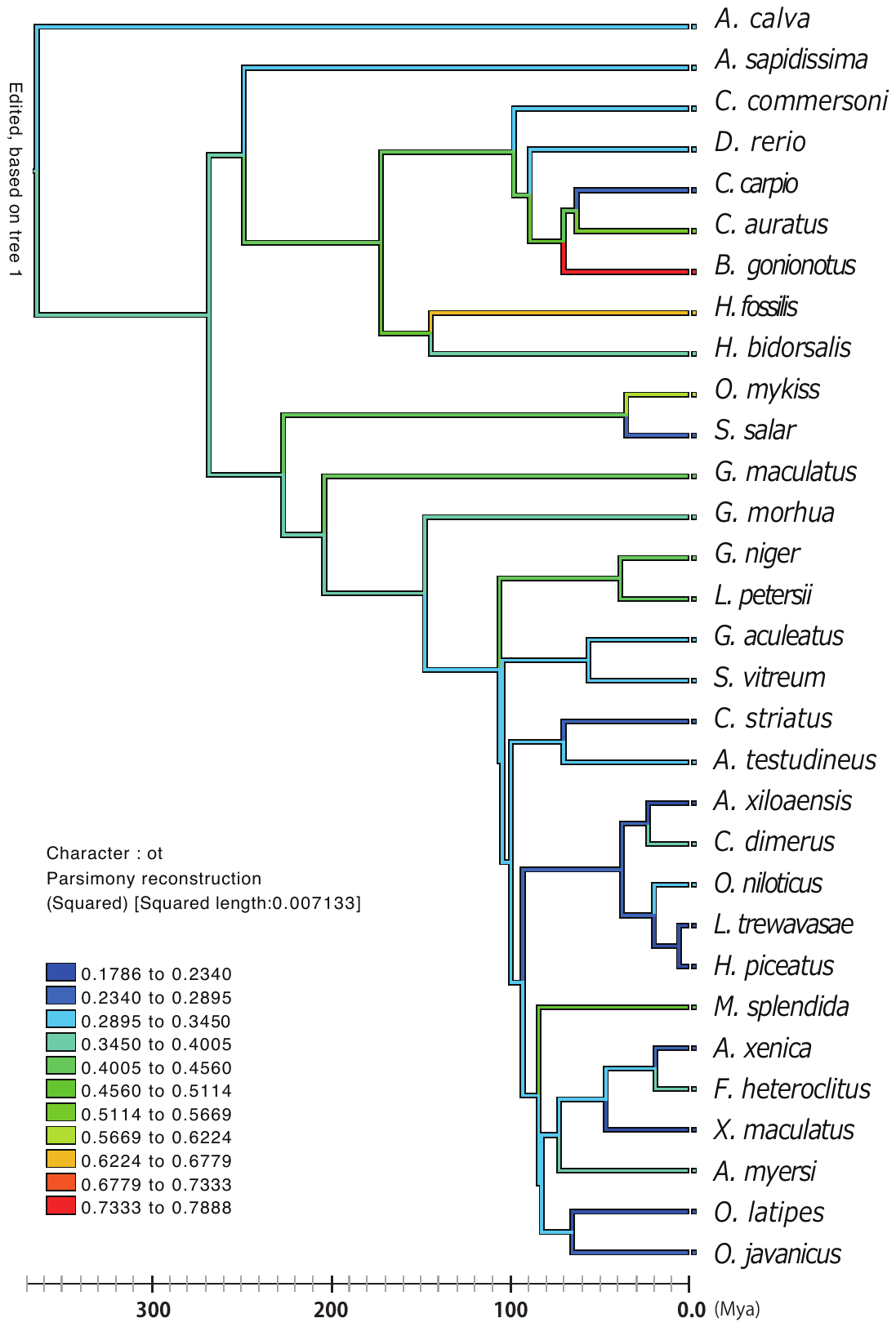

**Figure S7 Reconstructed ancestral ranks of otic vesicles/placodes/primordia (ot) by continuous analysis**

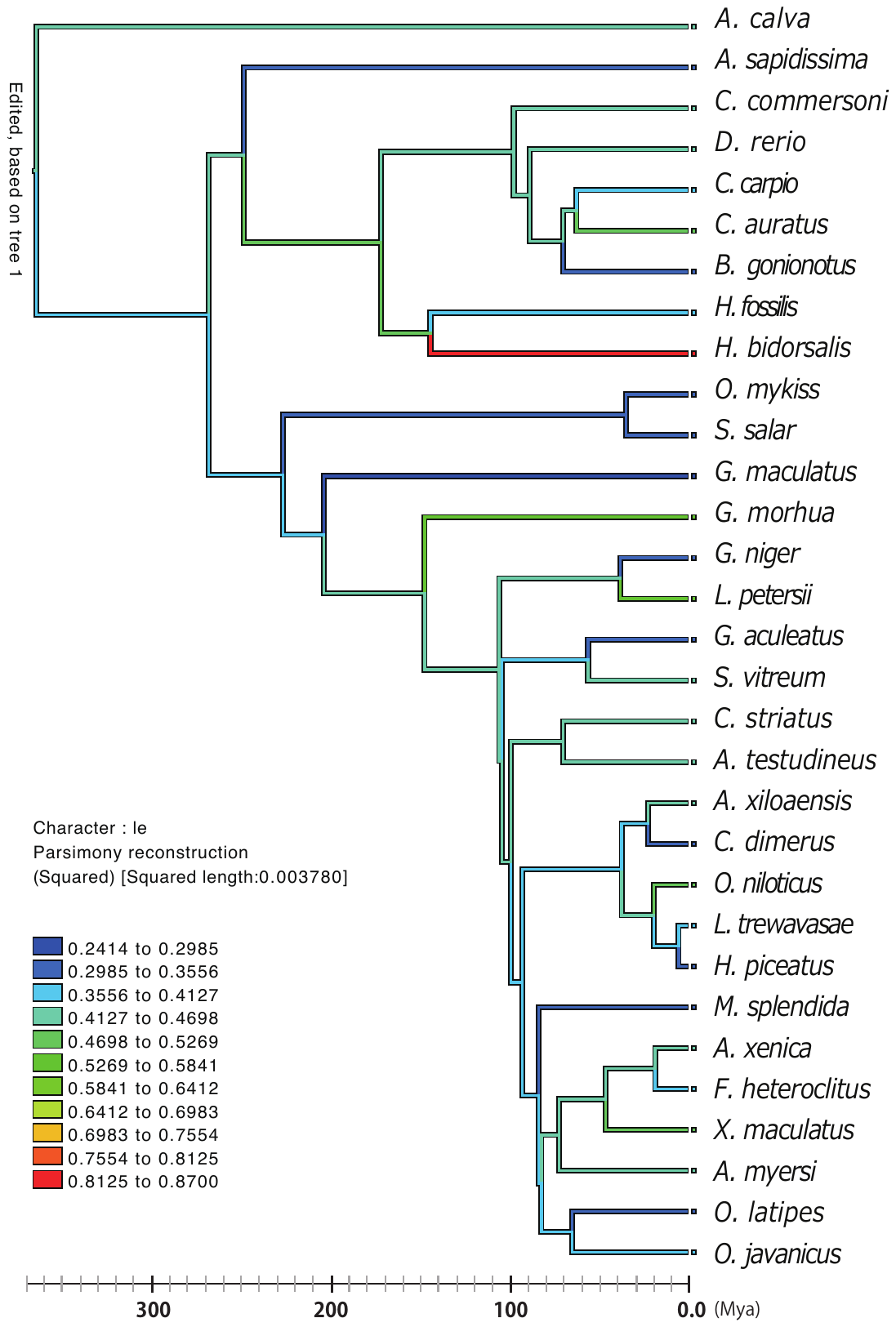

**Figure S8 Reconstructed ancestral ranks of lenses or lens placodes (le) by continuous analysis**

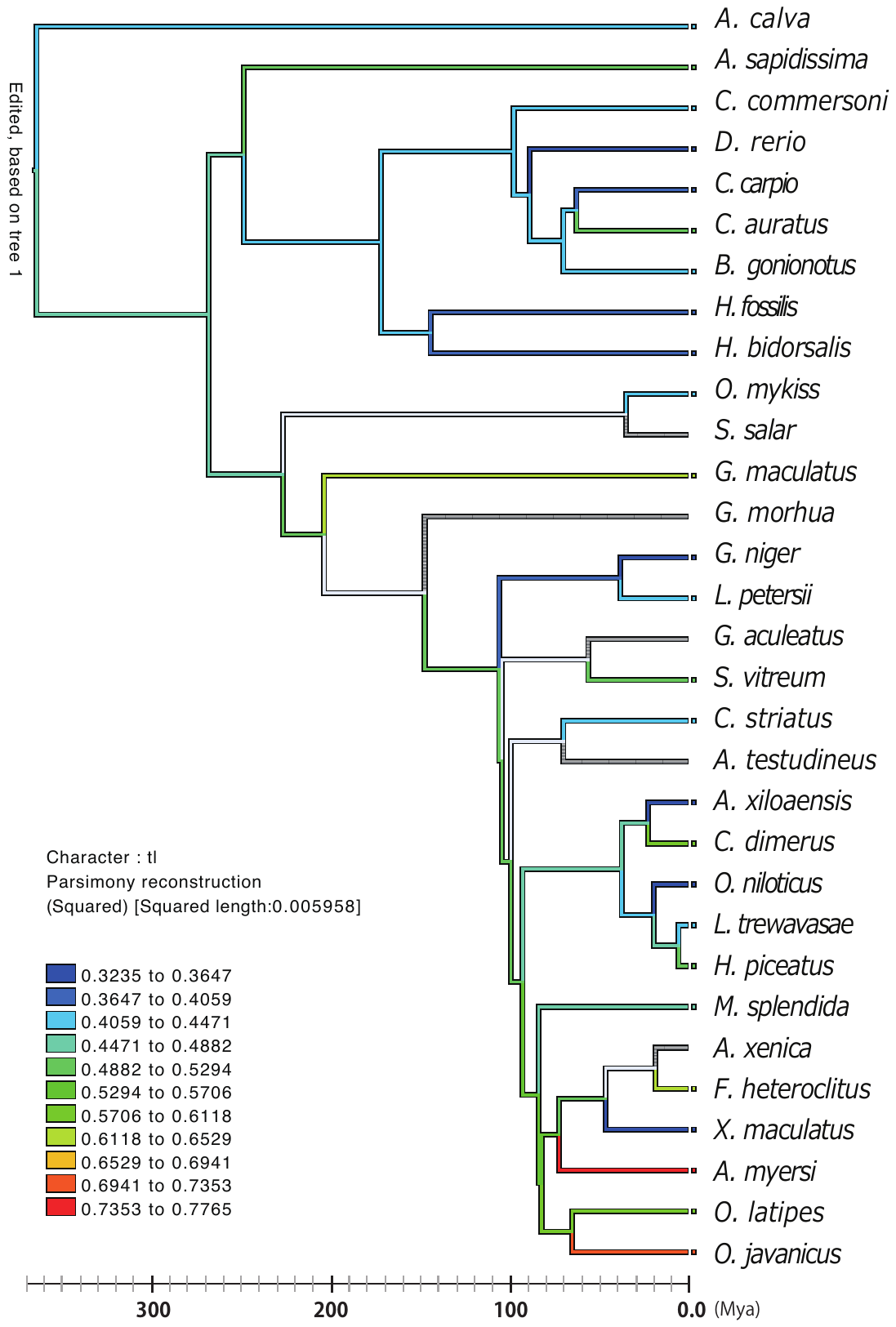

**Figure S9 Reconstructed ancestral ranks of tail lift from the yolk (tl) by continuous analysis**

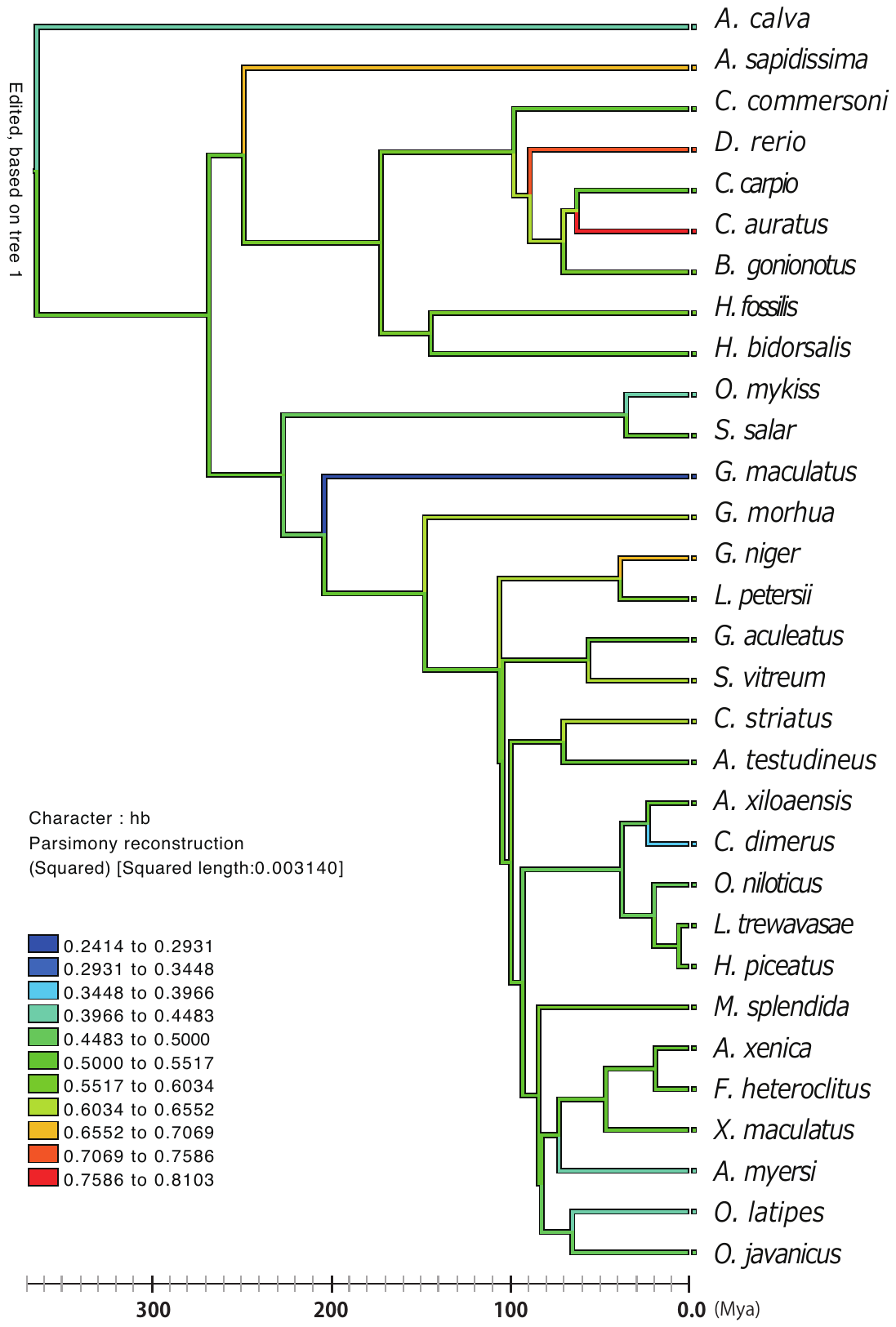

**Figure S10 Reconstructed ancestral ranks of heart beating/pulsing (hb) by continuous analysis**

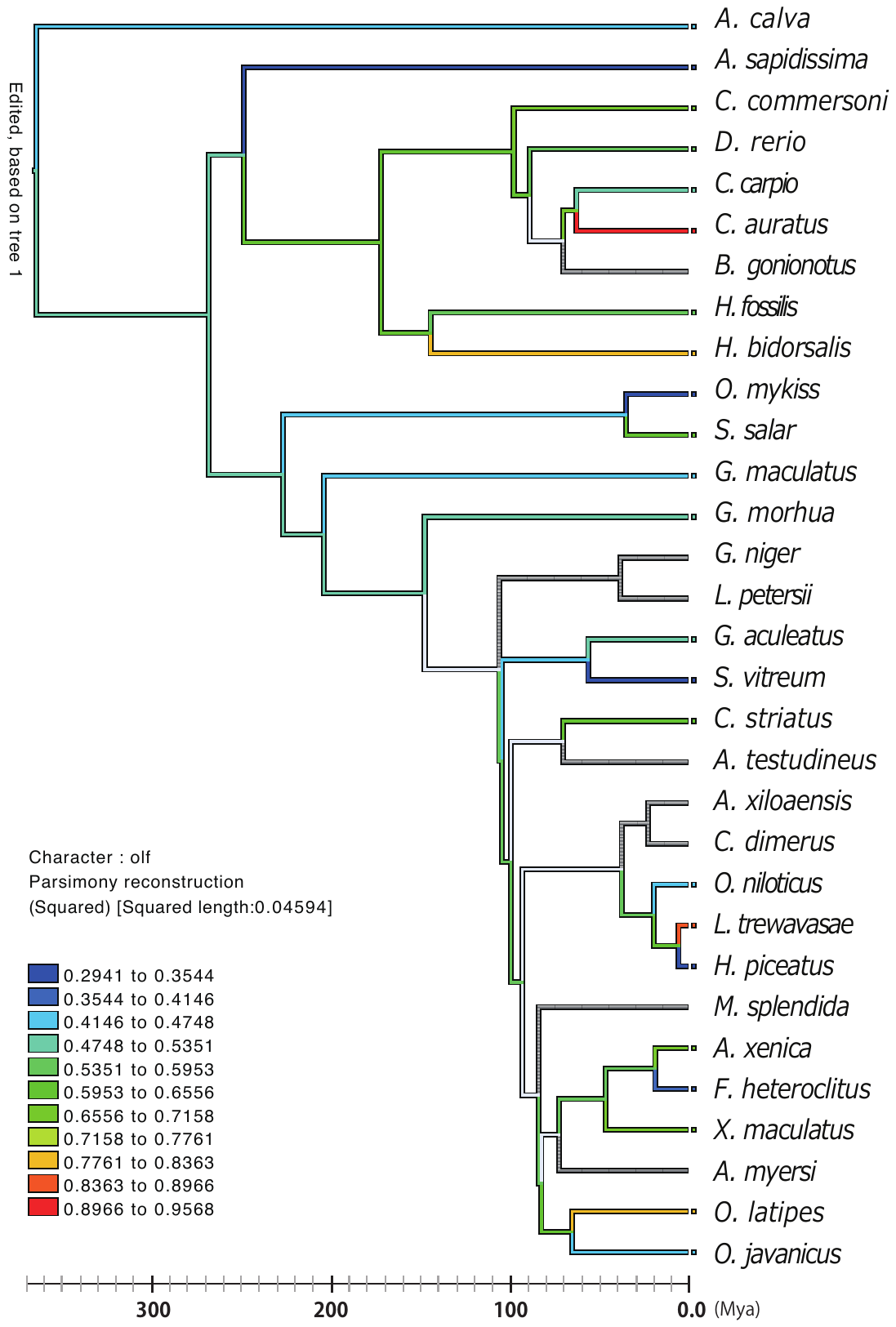

**Figure S11 Reconstructed ancestral ranks of olfactory vesicles/pits/placodes (olf) by continuous analysis**

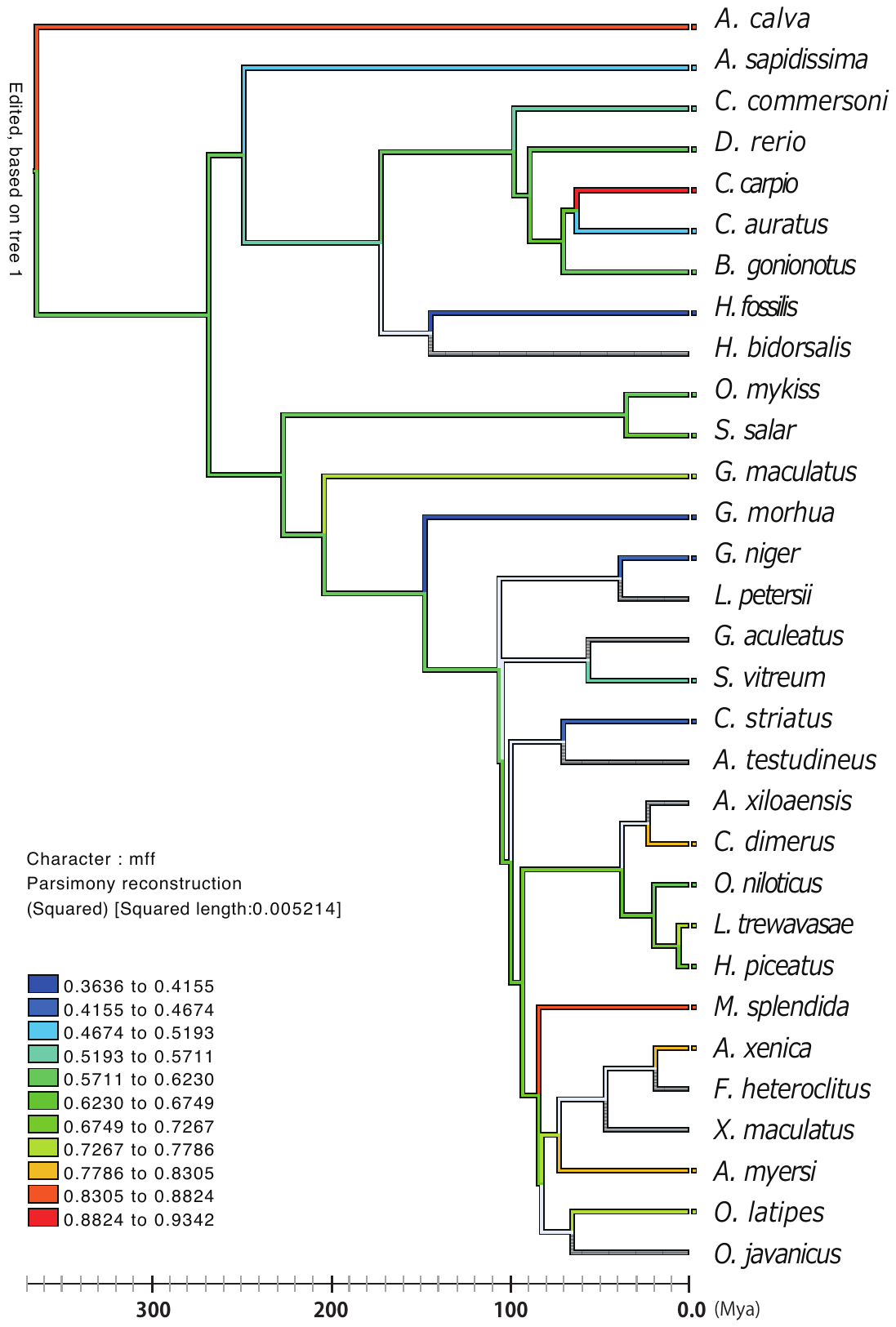

**Figure S12 Reconstructed ancestral ranks of a medial finfold (mff) by continuous analysis**

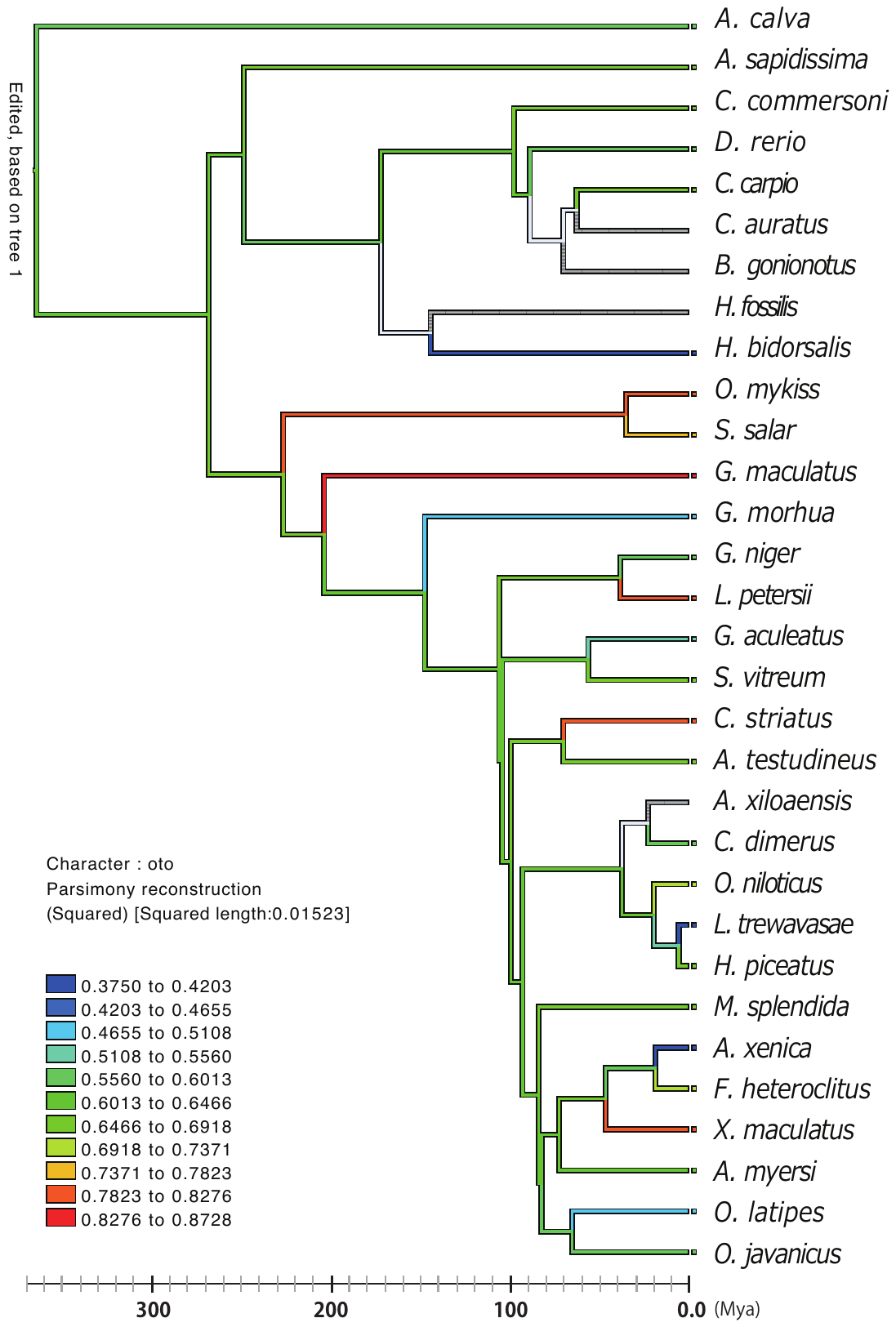

**Figure S13 Reconstructed ancestral ranks of otoliths (oto) by continuous analysis**

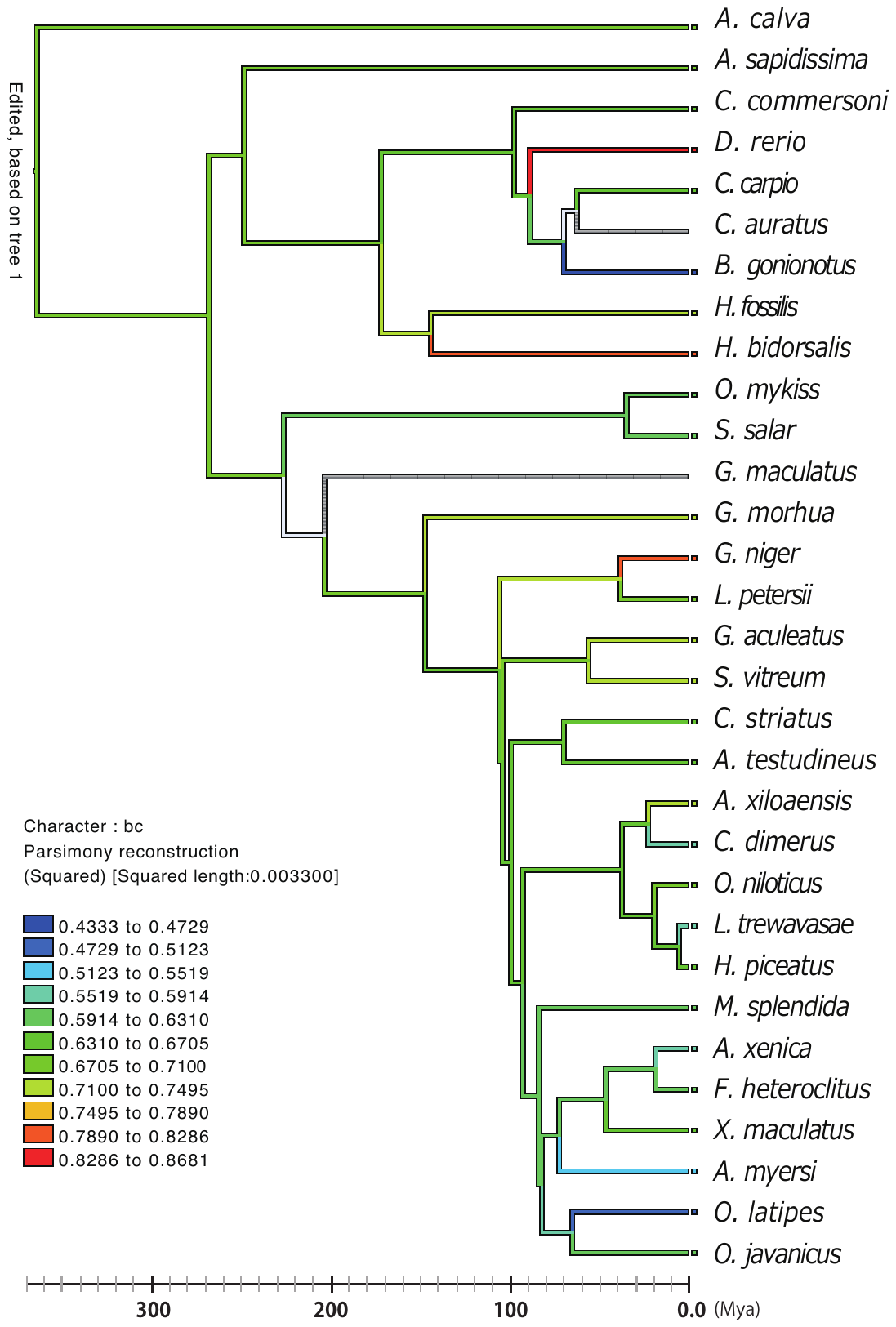

**Figure S14 Reconstructed ancestral ranks of blood circulation (bc) by continuous analysis**

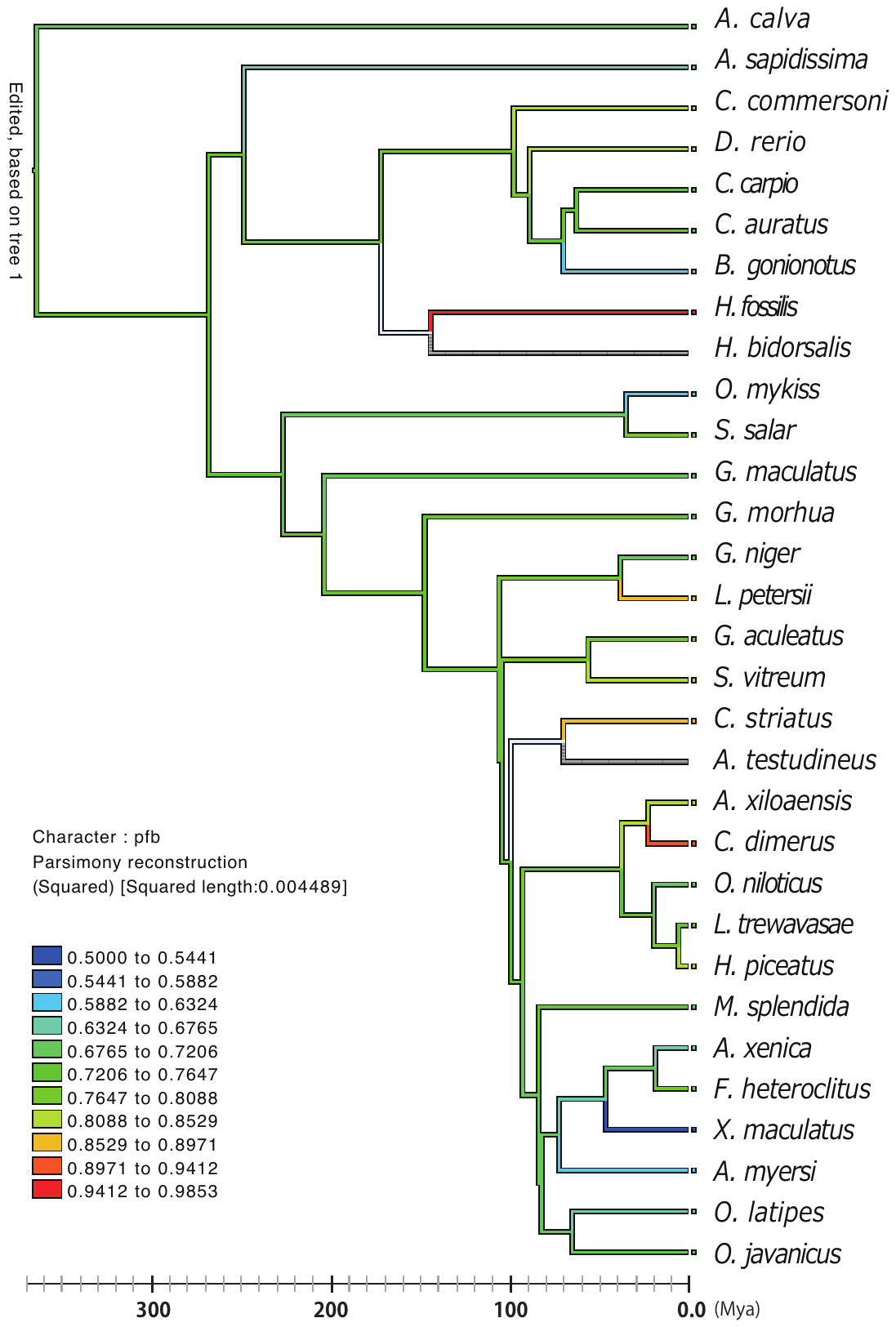

**Figure S15 Reconstructed ancestral ranks of pectoral fin buds (pfb) by continuous analysis**

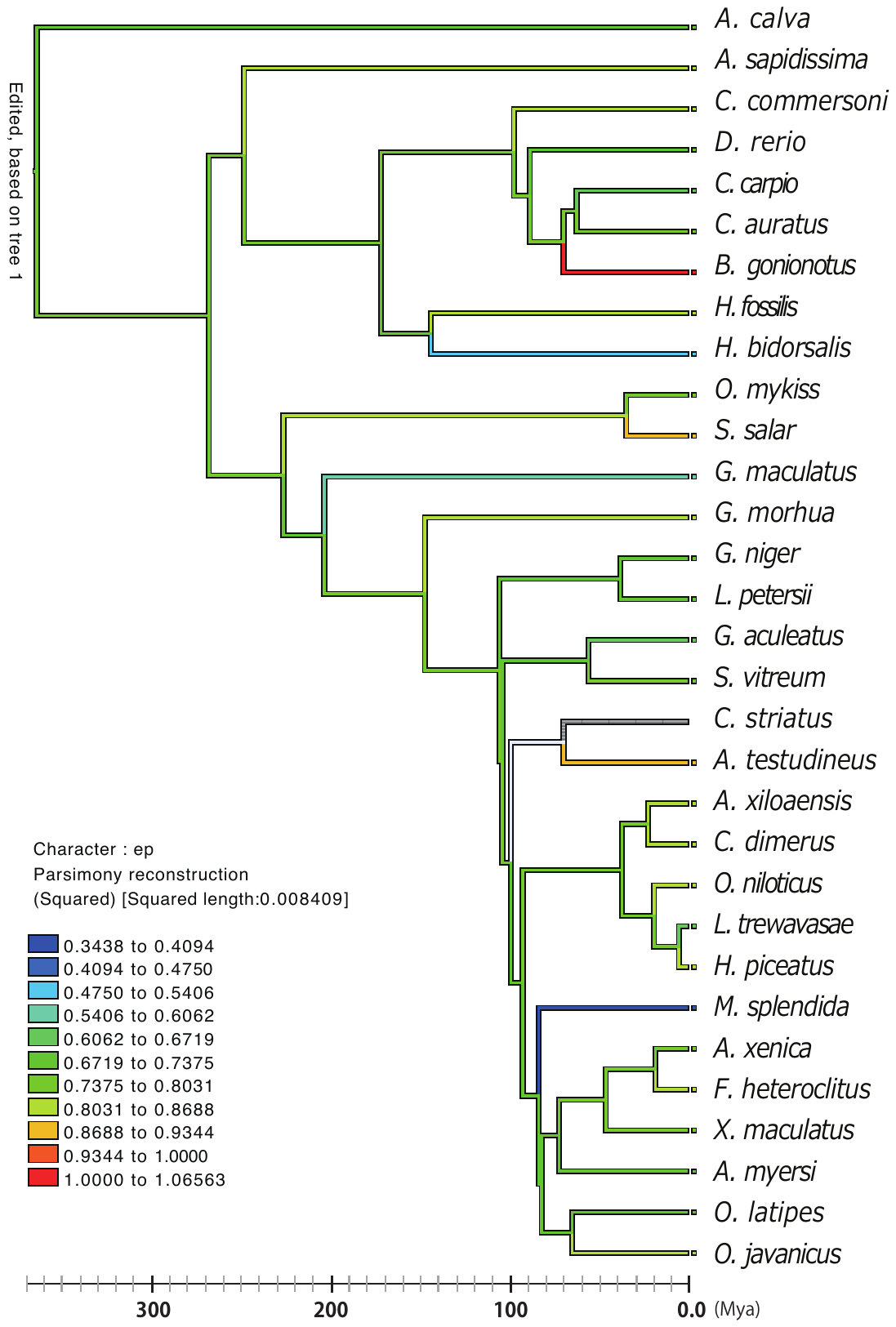

**Figure S16 Reconstructed ancestral ranks of eye pigmentation (ep) by continuous analysis**

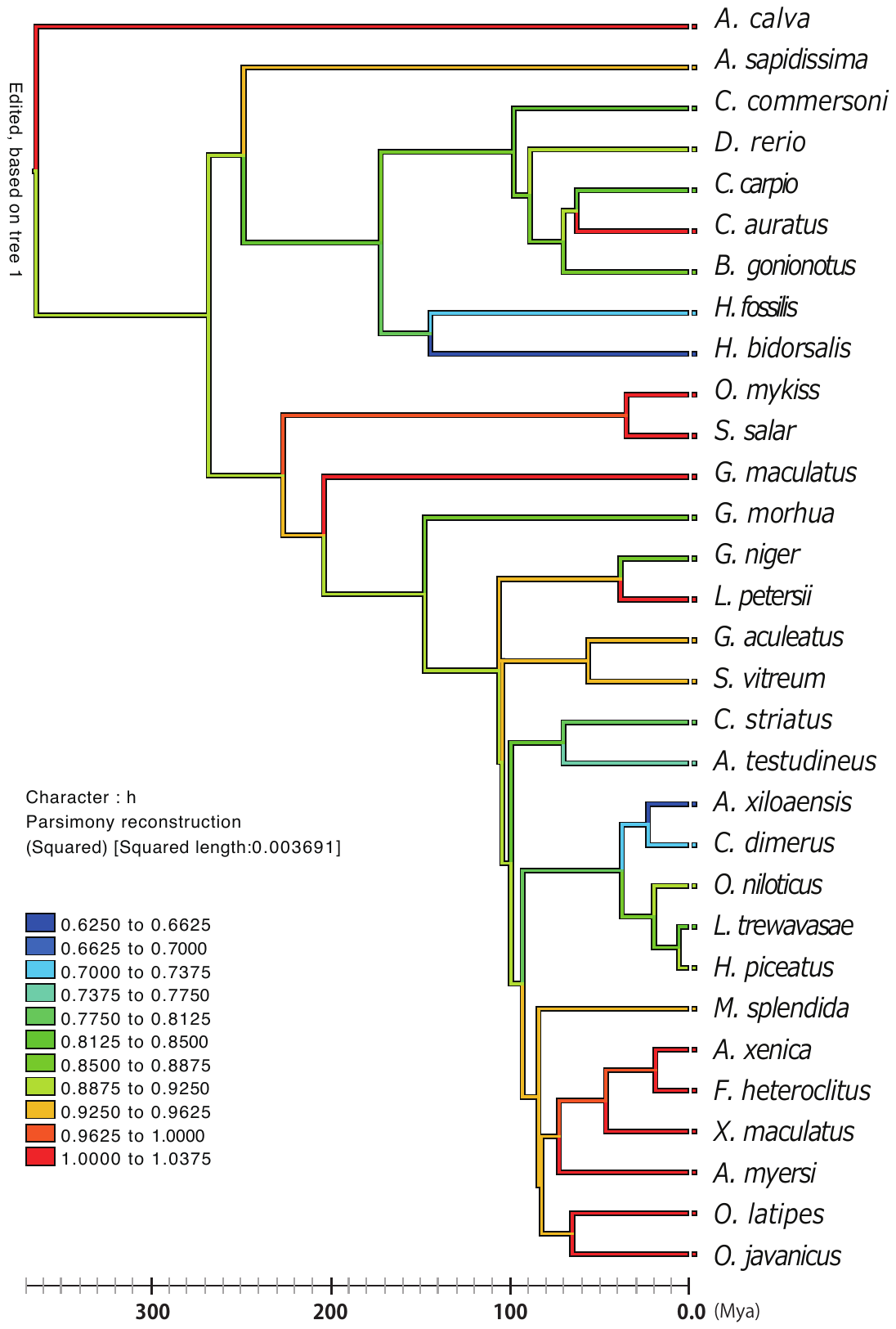

**Figure S17 Reconstructed ancestral ranks of hatching (h) by continuous analysis**

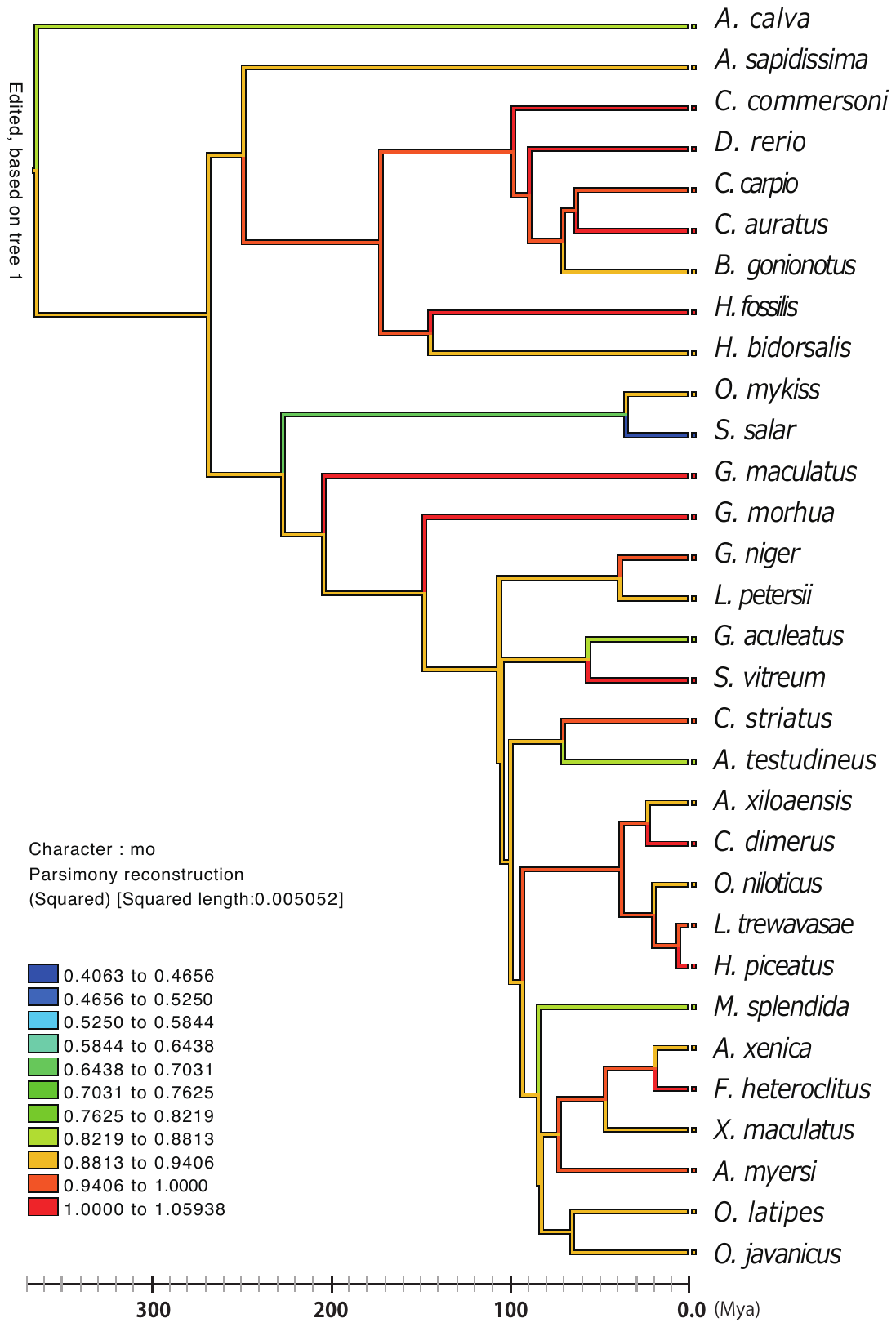

**Figure S18 Reconstructed ancestral ranks of a mouth opening (mo) by continuous analysis**

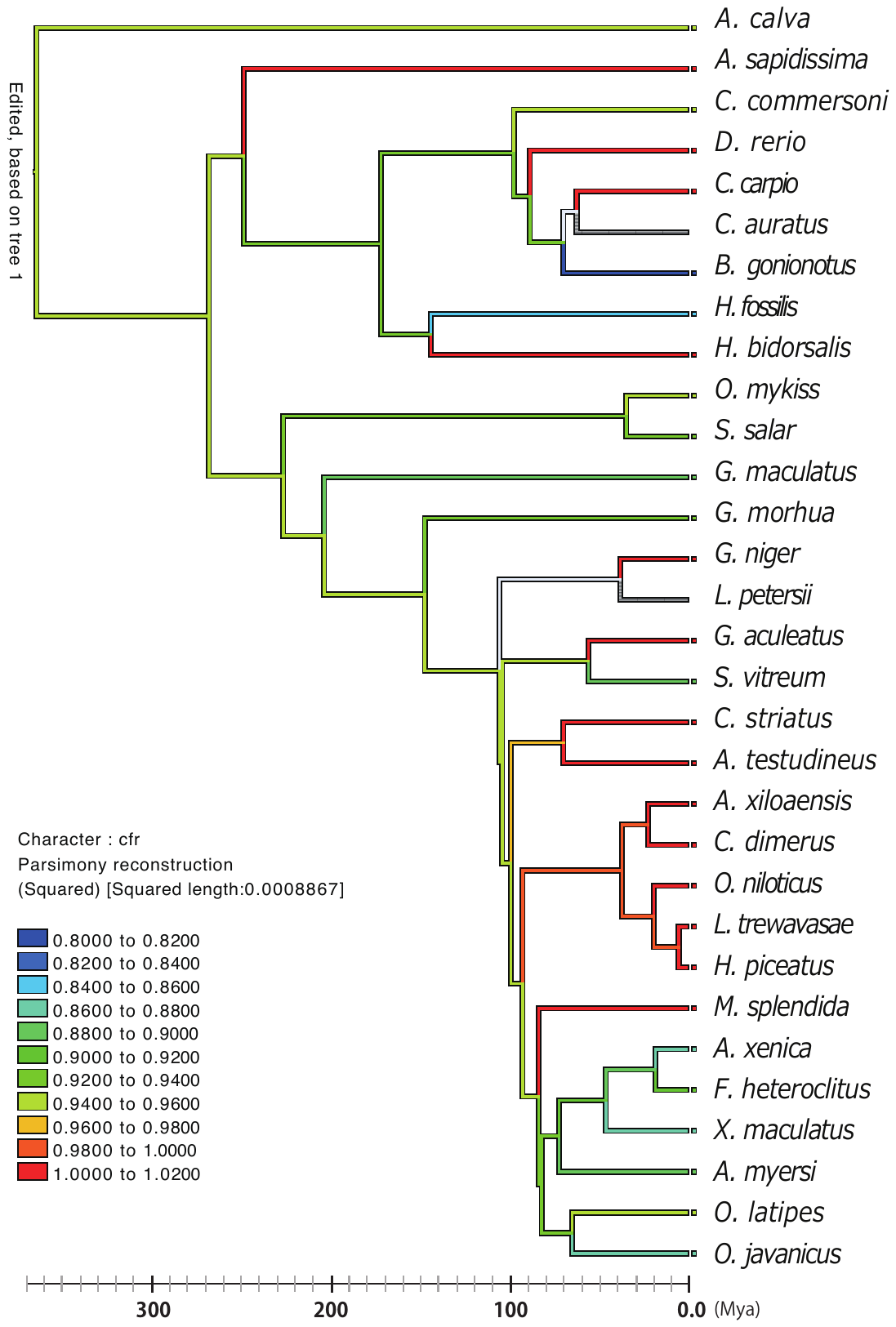

**Figure S19 Reconstructed ancestral ranks of caudal fin rays (cfr) by continuous analysis**

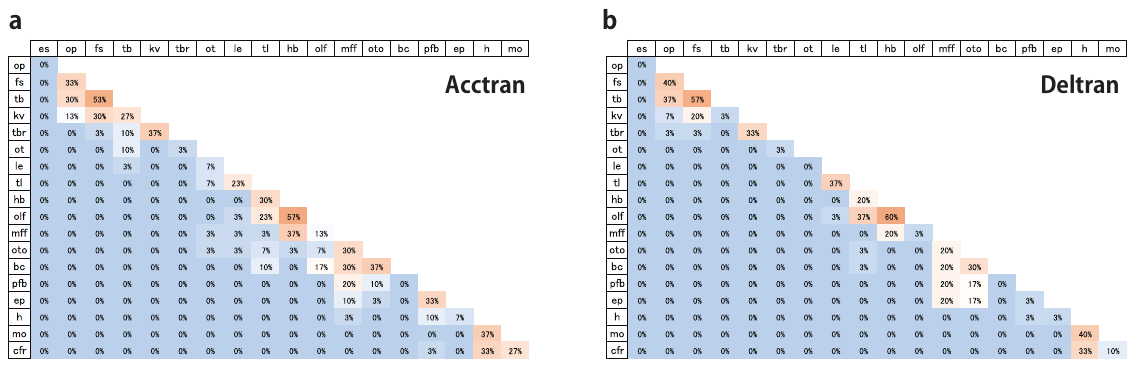

**Figure S20 Sequence orders of event pairs in ancestral developmental sequences**

The percentage of the sequences in which the row event occurs later than the column event calculated from the ancestral developmental sequences estimated by the event-pairing method under acctran (a) and deltran (b) optimizations.

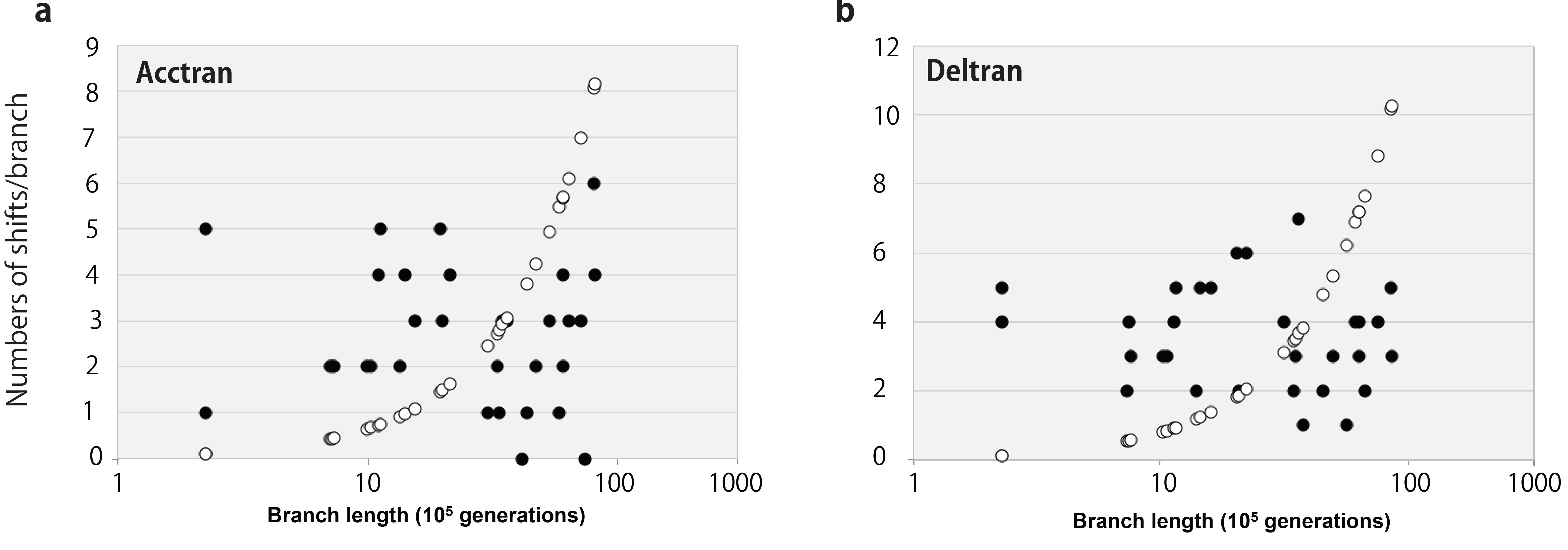

**Figure S21 Distribution of heterochronic shifts on the branches scaled by the generation number**

The relationships between the generation number and the number of heterochronic shifts estimated by the event-pairing method under acctran (a) and deltran (b) optimizations (black circle). Open circles show the distribution of reference data. The horizontal axis is scaled by the generation number considering the average generation time of each species (supplemental file5).

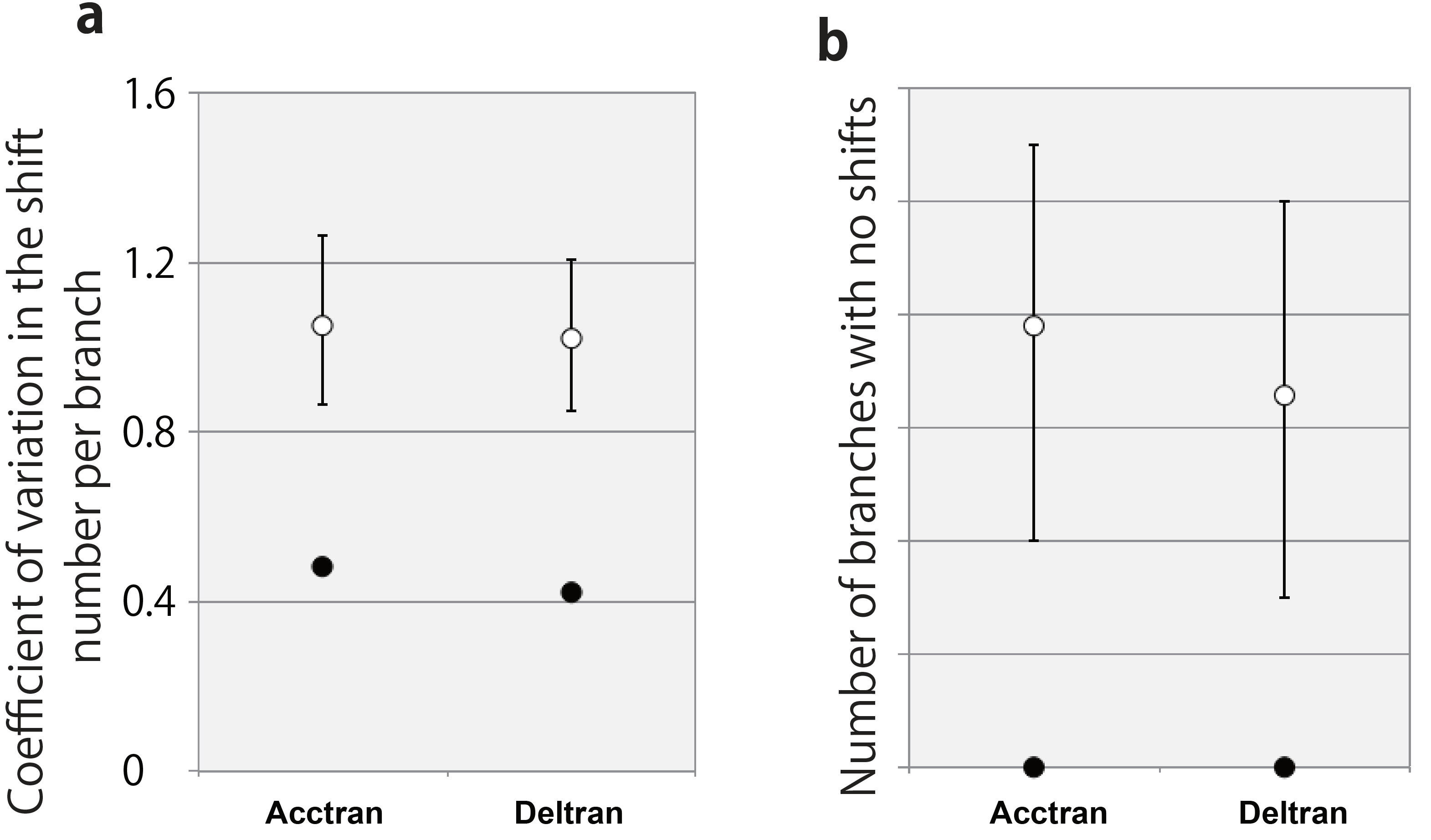

**Figure S22 Statistical comparisons of heterochronic shifts on the branches scaled by the generation number**

(a) The coefficient of variation for the number of heterochronic shifts in each branch (Figure 8c) rescaled by the generation number as phylogenetic time. (b) The number of branches with no heterochronic shifts (Figure 6d) rescaled by the generation number as phylogenetic time. The labels and marks are the same as those in Figure 6c and 6d.
