## Supplemental file.3 for "Frequent Non-random Shifts in the Temporal Sequence of Developmental Landmark Events during Teleost Evolutionary Diversification"

**Supplemental file 3 List of heterochronic shifts estimated by Parsimov**

The heterochronic shifts estimated in each branch are listed. E and L stand for earlier and later shifts, respectively. Twins indicate the rank change only between two events. Individual nodes in the phylogenetic tree are numbered as below.

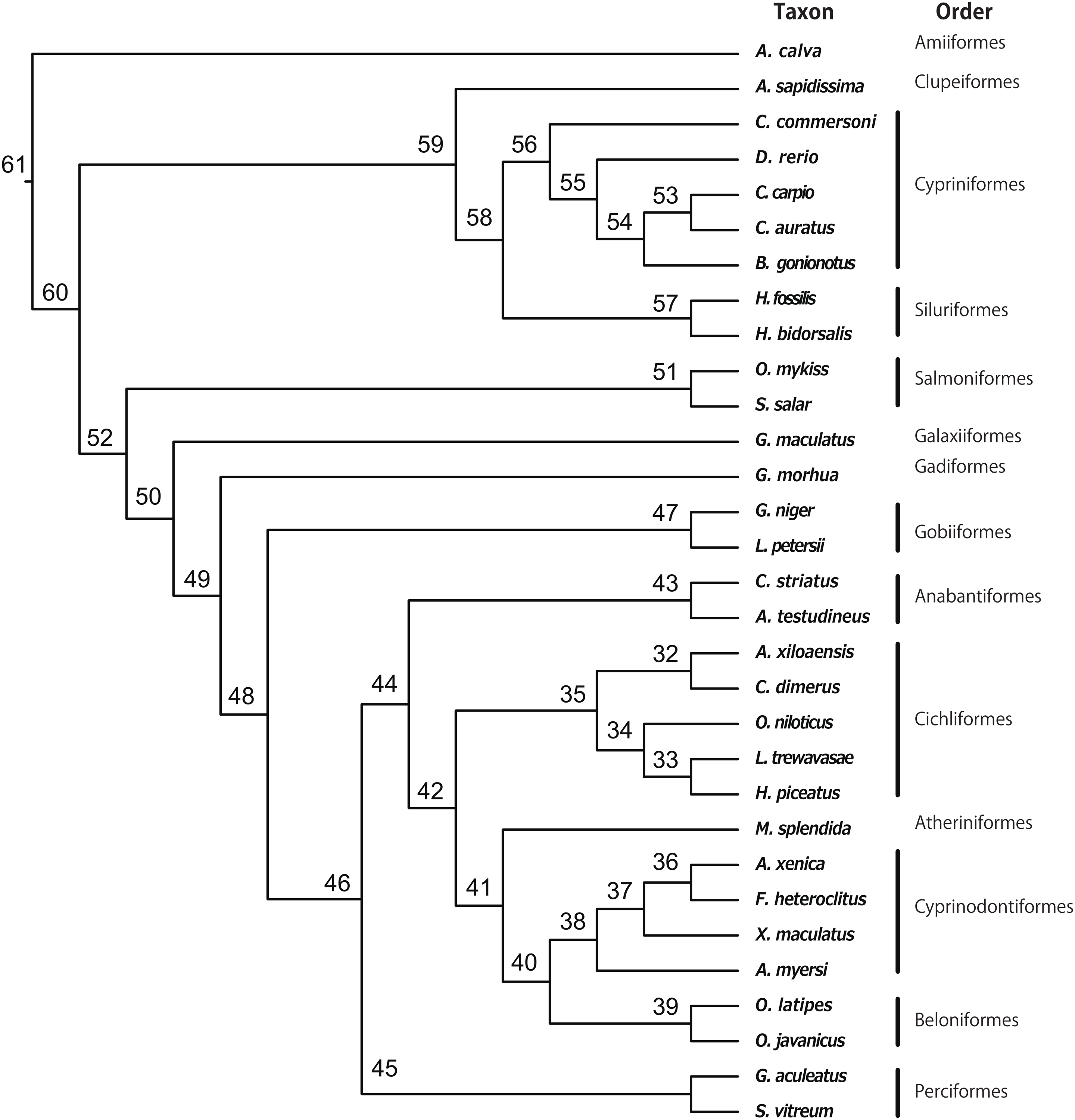

**Event number**

1: embryonic shield (es)

2: optic vesicles/placodes/primordia (op)

3: first somite (fs)

4: tail buds (tb)

5: Kupffer’s vesicle (kv)

6: three brain regionalization (tbr)

7: otic vesicles/placodes/primordia (ot)

8: lenses or lens placodes (le)

9: tail lift from the yolk (tl)

10: heart beating/pulsing (hb)

11: olfactory vesicles/pits/placodes (olf)

12: medial finfold (mff)

13: otoliths (oto)

14: blood circulation (bc)

15: pectoral fin buds (pfb)

16: eye pigmentation (ep)

17: hatching (h)

18: mouth opening (mo)

19: caudal fin rays (cfr)

**Acctram optimization**

======================================

Node 61 --> *A.calva*

Twins (9, 11)

======================================

Node 61 --> Node 60

Char 8 moved E relative to 9, 10, 11

Char 13 moved E relative to 14, 15, 18

======================================

Node 52 --> Node 50

Twins (2, 3) (13, 14)

Char 18 moved L relative to 17, 19

======================================

Node 50 --> Node 49

Twins (2, 6) (13, 15)

Char 9 moved E relative to 8, 10

Char 12 moved E relative to 7, 10

Char 17 moved E relative to 18, 19

======================================

Node 49 --> Node 48

Twins (14, 15) (18, 19)

Char 6 moved L relative to 4, 5, 7

======================================

Node 48 --> Node 46

Twins (2, 3)

======================================

Node 46 --> Node 44

Twins (5, 3) (16, 12)

======================================

Node 44 --> Node 42

Twins (5, 4)

Char 12 moved L relative to 9, 10, 13, 14

======================================

Node 42 --> Node 35

Twins (3, 2)

Char 11 moved E relative to 8, 14

======================================

Node 35 --> Node 32

Twins (13, 9)

Char 6 moved L relative to 8, 10

Char 17 moved E relative to 12, 15, 16

======================================

Node 32 --> *A.xiloaensis*

Twins (17, 14)

Char 7 moved E relative to 2, 3, 4

======================================

Node 32 --> *C.dimerus*

Twins (16, 15) (19, 18)

======================================

Node 35 --> Node 34

Twins (3, 4) (6, 7)

Char 12 moved E relative to 13, 14, 16

======================================

Node 34 --> Node 33

Char 4 moved L relative to 2, 6, 7

======================================

Node 33 --> *H.piceatus*

Twins (19, 18)

======================================

Node 33 --> *L.trewavasae*

Char 3 moved L relative to 2, 4, 7

Char 6 moved E relative to 2, 4

Char 11 moved L relative to 8, 10, 12, 14, 15, 17

Char 12 moved L relative to 14, 15

Char 13 moved E relative to 8, 9, 10, 14

======================================

Node 34 --> *O.niloticus*

Twins (3, 2) (18, 17)

Char 12 moved E relative to 13, 14

Char 15 moved E relative to 13, 14, 16

======================================

Node 42 --> Node 41

Twins (8, 9) (14, 13) (18, 17)

Char 11 moved L relative to 10, 15

Char 12 moved L relative to 15, 16

======================================

Node 41 --> Node 40

Twins (5, 2) (15, 16)

Char 9 moved L relative to 10, 13, 14

Char 19 moved E relative to 17, 18

======================================

Node 40 --> Node 38

Twins (3, 2) (6, 7) (10, 8) (15, 13)

======================================

Node 38 --> *A.myersi*

Char 2 moved L relative to 3, 4

Char 9 moved L relative to 15, 16

======================================

Node 38 --> Node 37

Twins (12, 16)

Char 4 moved L relative to 3, 6, 7

Char 5 moved L relative to 2, 6

Char 9 moved E relative to 13, 14

======================================

Node 37 --> Node 36

Twins (8, 10) (11, 9) (13, 15)

Char 3 moved L relative to 2, 6

======================================

Node 36 --> *F.heteroclitus*

Twins (17, 18)

Char 11 moved E relative to 8, 10, 14, 15

======================================

Node 36 --> *A.xenia*

Char 13 moved E relative to 8, 10, 14

======================================

Node 37 --> *X.maculatus*

Twins (7, 6)

Char 9 moved E relative to 8, 10, 14

Char 15 moved E relative to 8, 10, 11, 14

======================================

Node 40 --> Node 39

Twins (12, 11) (13, 14)

======================================

Node 39 --> *O.javanicus*

Char 11 moved E relative to 9, 10, 14, 15

======================================

Node 39 --> *O.latipes*

Twins (18, 19)

Char 7 moved E relative to 3, 6

Char 11 moved L relative to 13, 14, 15, 16

======================================

Node 41 --> *M.splendida*

Twins (18, 12)

Char 7 moved L relative to 8, 9

Char 16 moved E relative to 8, 9, 13, 14, 15

======================================

Node 44 --> Node 43

Char 12 moved E relative to 8, 11

======================================

Node 43 --> *C.striatus*

Char 5 moved L relative to 6, 7, 8

======================================

Node 43 --> *A.testudineus*

Twins (5, 2)

Char 3 moved L relative to 6, 7

======================================

Node 46 --> Node 45

Twins (4, 3) (8, 9) (16, 15)

======================================

Node 45 --> *G.aculeatus*

Twins (16, 14) (18, 17)

Char 5 moved L relative to 3, 7

Char 6 moved E relative to 2, 3

Char 13 moved E relative to 10, 11

======================================

Node 45 --> *S.vitreum*

Char 11 moved E relative to 6, 7, 8, 10

Char 19 moved E relative to 17, 18

======================================

Node 48 --> Node 47

Twins (9, 7) (12, 6) (16, 14)

======================================

Node 47 --> *G.niger*

Char 4 moved L relative to 2, 5

Char 6 moved L relative to 7, 8, 9

Char 7 moved L relative to 8, 9

Char 10 moved L relative to 13, 15, 16

Char 14 moved L relative to 15, 16

======================================

Node 47 --> *L.petersii*

Twins (6, 7) (9, 8) (18, 17)

======================================

Node 49 --> *G.morhua*

Twins (15, 16)

Char 3 moved E relative to 1, 2, 5

Char 4 moved L relative to 2, 5

======================================

Node 50 --> *G.maculatus*

Char 3 moved L relative to 2, 4, 6, 8

Char 5 moved L relative to 6, 8

Char 7 moved L relative to 8, 11

Char 10 moved E relative to 8, 9, 11

Char 15 moved E relative to 12, 13

Char 16 moved E relative to 9, 12, 13, 15

======================================

Node 52 --> Node 51

Twins (15, 16)

Char 4 moved L relative to 2, 3, 7, 8, 18

Char 5 moved E relative to 2, 6

Char 14 moved E relative to 12, 13

======================================

Node 51 --> *O.mykiss*

Char 7 moved L relative to 8, 10, 12, 14

Char 11 moved E relative to 8, 10

======================================

Node 51 --> *S.salar*

Char 11 moved L relative to 10, 12

Char 18 moved E relative to 10, 12, 13, 14, 16

======================================

Node 60 --> Node 59

Twins (3, 5) (17, 19)

Char 10 moved L relative to 9, 13

======================================

Node 59 --> Node 58

Twins (4, 6)

Char 9 moved E relative to 7, 8

Char 11 moved L relative to 12, 13

Char 15 moved L relative to 13, 14, 17

Char 18 moved L relative to 17, 19

======================================

Node 56 --> Node 55

Twins (15, 17) (16, 14)

Char 4 moved E relative to 2, 3

Char 12 moved L relative to 11, 13

======================================

Node 55 --> Node 54

Char 7 moved L relative to 8, 12

Char 12 moved L relative to 13, 14

Char 13 moved L relative to 11, 16

======================================

Node 54 --> *B.gonionotus*

Twins (3, 4)

Char 7 moved L relative to 9, 10, 12, 15

Char 16 moved L relative to 15, 17, 18

Char 19 moved E relative to 17, 18

======================================

Node 54 --> Node 53

Twins (2, 3) (18, 19)

======================================

Node 53 --> *C.auratus*

Twins (2, 6) (18, 17)

Char 11 moved L relative to 10, 15

Char 12 moved E relative to 8, 9, 15

======================================

Node 53 --> *C.carpio*

Char 6 moved E relative to 2, 5

Char 7 moved E relative to 5, 8, 9

Char 12 moved L relative to 10, 11, 15, 16, 17

Char 15 moved L relative to 10, 16

======================================

Node 55 --> *D.rerio*

Char 6 moved L relative to 2, 7

Char 10 moved L relative to 11, 12, 13

Char 14 moved L relative to 13, 15

======================================

Node 56 --> *C.commersoni*

Twins (17, 16) (19, 18)

Char 11 moved L relative to 10, 14

======================================

Node 58 --> Node 57

Char 2 moved E relative to 3, 6

Char 12 moved E relative to 5, 7, 8

Char 13 moved E relative to 6, 7, 8, 9, 10, 11

Char 15 moved L relative to 16, 19

======================================

Node 57 --> *H.bidorsalis*

Twins (18, 19)

Char 6 moved L relative to 5, 7, 9, 10

Char 8 moved L relative to 9, 10, 11, 14

Char 11 moved L relative to 10, 14

======================================

Node 57 --> *H.fossilis*

Char 5 moved L relative to 6, 8, 9

Char 7 moved L relative to 8, 9, 10, 11

======================================

Node 59 --> *A.sapidissima*

Char 4 moved L relative to 2, 3, 7

======================================

**Deltran optimization**

======================================

Node 61 --> *A.calva*

Twins (9, 11)

======================================

Node 61 --> Node 60

Twins (8, 10) (13, 18) (14, 16)

======================================

Node 52 --> Node 50

Twins (17, 18)

======================================

Node 50 --> Node 49

Twins (2, 6) (12, 10)

Char 13 moved E relative to 14, 15, 16

Char 17 moved E relative to 18, 19

======================================

Node 49 --> Node 48

Twins (2, 3) (9, 10)

Char 6 moved L relative to 4, 5

======================================

Node 48 --> Node 46

Twins (14, 15)

======================================

Node 46 --> Node 44

Twins (9, 11)

======================================

Node 44 --> Node 42

Char 12 moved L relative to 10, 13, 14

======================================

Node 35 --> Node 32

Char 6 moved L relative to 7, 8, 10

Char 17 moved E relative to 15, 16

======================================

Node 32 --> *A.xiloaensis*

Twins (17, 14)

Char 7 moved E relative to 2, 3, 4

======================================

Node 32 --> *C.dimerus*

Twins (19, 18)

Char 9 moved L relative to 13, 14

Char 12 moved L relative to 13, 14, 17

======================================

Node 34 --> Node 33

Char 4 moved L relative to 2, 6, 7

======================================

Node 33 --> *H.piceatus*

Twins (19, 18)

Char 4 moved L relative to 3, 7, 8

Char 6 moved L relative to 7, 8

Char 9 moved L relative to 8, 10

======================================

Node 33 --> *L.trewavasae*

Char 3 moved L relative to 2, 4, 7

Char 6 moved E relative to 2, 4

Char 11 moved L relative to 8, 10, 12, 14, 15, 17

Char 12 moved L relative to 14, 15

Char 13 moved E relative to 8, 9, 10, 14

======================================

Node 34 --> *O.niloticus*

Twins (18, 17)

Char 3 moved E relative to 2, 4

Char 12 moved E relative to 13, 14

Char 15 moved E relative to 13, 14, 16

======================================

Node 42 --> Node 41

Twins (8, 9) (18, 17)

Char 5 moved E relative to 3, 4

Char 12 moved L relative to 13, 14, 15, 16

======================================

Node 41 --> Node 40

Twins (8, 11) (15, 16) (19, 17)

Char 9 moved L relative to 10, 14

======================================

Node 40 --> Node 38

Twins (14, 13) (19, 18)

======================================

Node 38 --> *A.myersi*

Twins (10, 8)

Char 2 moved L relative to 3, 5

Char 9 moved L relative to 13, 14, 16

======================================

Node 36 --> *F.heteroclitus*

Twins (17, 18)

Char 11 moved E relative to 8, 9, 10

======================================

Node 36 --> *A.xenia*

Twins (12, 16)

Char 11 moved L relative to 10, 14, 15

Char 13 moved E relative to 8, 10, 14

======================================

Node 37 --> *X.maculatus*

Char 4 moved L relative to 2, 3, 6

======================================

Node 40 --> Node 39

Twins (7, 6)

Char 9 moved L relative to 13, 14

======================================

Node 39 --> *O.javanicus*

Twins (19, 18)

Char 9 moved L relative to 11, 15

======================================

Node 39 --> *O.latipes*

Char 7 moved E relative to 3, 6

Char 11 moved L relative to 10, 12, 13, 14, 15, 16

======================================

Node 41 --> *M.splendida*

Twins (18, 12)

Char 7 moved L relative to 8, 9

Char 14 moved E relative to 10, 13

Char 16 moved E relative to 8, 9, 10, 13, 14, 15

======================================

Node 44 --> Node 43

Twins (7, 6)

======================================

Node 43 --> *C.striatus*

Char 5 moved L relative to 3, 6, 7, 9

Char 12 moved E relative to 8, 9

Char 14 moved E relative to 10, 13

Char 17 moved E relative to 13, 15

======================================

Node 43 --> *A.testudineus*

Char 3 moved L relative to 2, 6, 7

Char 16 moved L relative to 17, 18

======================================

Node 46 --> Node 45

Twins (2, 3) (13, 14) (16, 15)

======================================

Node 45 --> *G.aculeatus*

Twins (16, 14) (18, 17)

Char 5 moved L relative to 3, 7

Char 6 moved E relative to 2, 3

Char 8 moved E relative to 7, 11

Char 13 moved E relative to 10, 11

======================================

Node 45 --> *S.vitreum*

Twins (4, 3)

Char 9 moved L relative to 8, 12

Char 11 moved E relative to 7, 8, 10

Char 19 moved E relative to 17, 18

======================================

Node 48 --> Node 47

Twins (9, 7) (16, 14)

======================================

Node 47 --> *G.niger*

Char 4 moved L relative to 2, 5

Char 6 moved L relative to 7, 8, 9, 12

Char 7 moved L relative to 8, 9, 12

Char 10 moved L relative to 13, 15, 16

Char 14 moved L relative to 13, 15, 16

======================================

Node 47 --> *L.petersii*

Twins (9, 8) (18, 17)

======================================

Node 49 --> *G.morhua*

Twins (15, 16) (19, 18)

Char 3 moved E relative to 1, 5

Char 4 moved L relative to 2, 5

Char 11 moved E relative to 8, 10

Char 12 moved E relative to 7, 8, 11

Char 13 moved E relative to 8, 10, 11, 14

======================================

Node 50 --> *G.maculatus*

Twins (19, 18)

Char 3 moved L relative to 2, 4

Char 8 moved E relative to 9, 11

Char 10 moved E relative to 8, 9, 11

Char 16 moved E relative to 9, 15

======================================

Node 52 --> Node 51

Twins (5, 2) (14, 12) (15, 16)

======================================

Node 51 --> *O.mykiss*

Twins (5, 6)

Char 7 moved L relative to 9, 10, 12, 14

Char 11 moved E relative to 8, 10

======================================

Node 51 --> *S.salar*

Char 4 moved L relative to 2, 3, 5, 7, 8

Char 11 moved L relative to 8, 10, 12

Char 18 moved E relative to 10, 12, 13, 14

======================================

Node 60 --> Node 59

Twins (17, 19)

Char 10 moved L relative to 9, 12

Char 13 moved E relative to 14, 15, 16

======================================

Node 59 --> Node 58

Twins (3, 5) (14, 15) (17, 18)

Char 11 moved L relative to 9, 12, 13

======================================

Node 58 --> Node 56

Twins (8, 11) (13, 15) (19, 18)

======================================

Node 56 --> Node 55

Twins (9, 7) (13, 12)

Char 4 moved E relative to 2, 6

======================================

Node 55 --> Node 54

Twins (14, 12)

======================================

Node 54 --> *B.gonionotus*

Char 7 moved L relative to 9, 10, 12

Char 16 moved L relative to 17, 18

Char 19 moved E relative to 17, 18

======================================

Node 54 --> Node 53

Twins (2, 3)

======================================

Node 53 --> *C.auratus*

Twins (2, 6) (4, 3) (18, 17)

Char 10 moved L relative to 15, 16

Char 11 moved L relative to 10, 15, 16

Char 12 moved E relative to 8, 9

======================================

Node 53 --> *C.carpio*

Twins (18, 19)

Char 6 moved E relative to 2, 5

Char 7 moved E relative to 5, 9

Char 12 moved L relative to 10, 13, 15, 17

Char 16 moved E relative to 14, 15

======================================

Node 55 --> *D.rerio*

Twins (4, 3)

Char 6 moved L relative to 2, 7

Char 10 moved L relative to 11, 12, 13

Char 14 moved L relative to 13, 15

======================================

Node 56 --> *C.commersoni*

Twins (3, 4) (19, 18)

Char 5 moved L relative to 6, 7

Char 11 moved L relative to 10, 12, 14

Char 17 moved E relative to 15, 16

======================================

Node 58 --> Node 57

Twins (9, 7) (17, 14)

======================================

Node 57 --> *H.bidorsalis*

Char 6 moved L relative to 5, 7, 9, 10

Char 8 moved L relative to 9, 10, 14

Char 11 moved L relative to 10, 14

Char 13 moved E relative to 7, 9

======================================

Node 57 --> *H.fossilis*

Char 5 moved L relative to 6, 9

Char 7 moved L relative to 9, 10, 11

Char 12 moved E relative to 8, 9, 10

======================================

Node 59 --> *A.sapidissima*

Char 4 moved L relative to 2, 3, 7, 8, 11

Char 7 moved L relative to 8, 11

Char 9 moved L relative to 8, 12

Char 10 moved L relative to 11, 12, 13, 14

======================================
